# Stabilization of HSA32, an aggregation-prone protein, by the protein disaggregase HSP101 plays a critical role in maintaining acquired thermotolerance in Arabidopsis

**DOI:** 10.1101/2024.10.15.618377

**Authors:** Suma Mitra, Shih-Jiun Yu, Nai-Yu Liu, Chuan-Chih Hsu, Hong-Yi Li, Akankshita Borah, Yu-Yen Shen, Shang-hao Wu, Yang-Hsin Hsu, Hongyong Fu, Yee-yung Charng

**Affiliations:** Agricultural Biotechnology Research Center, Academia Sinica, Taipei, Taiwan, ROC; Molecular and Biological Agricultural Sciences program, Taiwan International Graduate Program, Academia Sinica, Taiwan, ROC; Graduate Institute of Biotechnology, National Chung Hsing University, Taichung, Taiwan, ROC; Department of Biochemical Science and Technology, National Taiwan University, Taipei, Taiwan, ROC; Institute of Plant and Microbial Biology, Academia Sinica, Taipei, Taiwan, ROC; Department of Horticulture, National Chiayi University, Chiayi, Taiwan, ROC

## Abstract

Heat acclimation confers acquired thermotolerance (AT), and heat-acclimation memory (HAM) is the maintenance of AT for some time. In Arabidopsis and rice, the heat-stress-associated 32-kDa protein (HSA32) and the protein disaggregase heat shock protein101 (HSP101) form a positive feedback loop at the protein level to maintain AT; HSA32 mitigates HSP101 degradation, and HSP101 positively regulates the accumulation of HSA32. Here, we report the underpinning mechanism regarding how HSP101 affects the HSA32 level in Arabidopsis. We found that, without HSP101, nascent HSA32 was rapidly degraded, and the proteasome inhibitor, bortezomib, inhibited the degradation. In response to heat stress, the nascent HSA32-GFP fusion protein was present in liquid condensates and diffused in the cytosol after returning to non-stress temperature. Proximity labeling with HSA32-TurboID identified HSP101 and five other protein chaperones and co-chaperones as the primary interactors. Disturbing the interaction between HSA32 and HSP101 destabilized HSA32 and compromised HAM. HSA32 is predicted as a TIM-barrel protein with three intrinsically disordered regions of high aggregation propensity. Recombinant HSA32 expressed in *E. coli* was partitioned into insoluble fractions, suggesting that HSA32 is aggregation-prone. Our findings highlight how the interplay between an aggregation-prone protein and a protein disaggregase can maintain plant stress memory.

## INTRODUCTION

Acclimation or priming is essential for plants to defend against abiotic and biotic stresses. This process prepares plants to better respond to a stressful challenge based on their prior experience of the same or similar situation. Recently, an emerging concept about acclimation/priming in plants is that memory is involved in this defense behavior without involving a nerve system (Wu et al., 2013; Kinoshita and Seki, 2014; Hilker et al., 2016; Martinez-Medina et al., 2016). Since memory affects behavior, knowing how plants retain memory is essential to better understanding their responses to diverse environmental cues. However, the mechanisms underlying the maintenance of plant stress memory are largely unclear. Recent progress in heat acclimation memory (HAM) provides examples of identifying components relevant to memory maintenance in plants (Charng et al., 2023).

When encountering elevated temperatures, plant cells respond promptly by producing a variety of protecting molecules, most prominently of the highly conserved heat-shock proteins (HSPs) families, which function as molecular chaperones in maintaining proteostasis (Hartl et al., 2011). There are five prominent families of HSPs usually classified and named according to their molecular weight: HSP100 (ClpB), HSP90, HSP70, the chaperonins (GroEL and HSP60), and the small HSP (sHSP) (Wang et al., 2004). The induction of certain HSP family members substantially increases the thermotolerance of plants to more severe heat encountered later on (Yeh et al., 2012), a process known as heat acclimation or thermal priming. The acquired thermotolerance (AT) contributed by the heat-induced HSPs decays when these HSPs gradually diminish after the plants return to non-stress conditions (Charng et al., 2023). AT is usually assessed by applying a heat stress (HS) regime consisting of two HS episodes, one being mild HS as an acclimation/priming treatment followed by much more severe triggering HS. The memory capacity is evaluated by introducing a recovery period of varied time between the two HS episodes. This recovery period is the memory phase (Hilker et al., 2016). AT serves as the proxy of the primed state that can be maintained for some time, and the retainment of AT is referred to as HAM (Wu et al., 2013; Charng et al., 2023).

Reverse and forward genetic studies on Arabidopsis have identified distinct components of HAM based on the assessment of AT after a short and a long memory phase, *i.e.*, short-term AT (SAT) and long-term AT (LAT), respectively (Yeh et al., 2012; Charng et al., 2023). Some component mutants show defective LAT but normal SAT, suggesting that these components are required for maintenance rather than the acquisition of AT (Charng et al., 2006; Charng et al., 2007; Wu et al., 2013; Stief et al., 2014; Brzezinka et al., 2016; Sedaghatmehr et al., 2016; Brzezinka et al., 2019; Urrea Castellanos et al., 2020; Friedrich et al., 2021; Crawford et al., 2024). Based on the known molecular functions of the identified components, HAM involves regulatory networks that modulate the stability of HS-induced HSPs, which are considered AT effectors (Charng et al., 2023). One of the best-characterized effectors is HSP101, a member of the HSP100 family, required for both SAT and LAT (Hong and Vierling, 2000; Queitsch et al., 2000; Wu et al., 2013). The orthologs of HSP101 in *E. coli* (ClpB) and yeast (HSP104) act as a protein disaggregase for solubilization and refolding of heat-denatured proteins in concert with other chaperones (Weibezahn et al., 2004; Sweeny and Shorter, 2016; Gates et al., 2017). Disaggregation and refolding of denatured proteins are essential for cell survival after severe HS (Hong et al., 2003; Weibezahn et al., 2004). In Arabidopsis, HSP101 is gradually degraded partly via the autophagy pathway, concomitant with the decay of AT during the memory phase (Wu et al., 2013; Sedaghatmehr et al., 2019). The stability of HSP101 is positively regulated by the heat-stress-associated 32-kDa protein (HSA32). Without HSA32, HSP101 is selectively degraded faster, leading to a faster decay of AT (Wu et al., 2013). The differential levels of HSP101 explain the phenotype of the *HSA32* knockout mutant, being defective in LAT but not SAT (Charng et al., 2006).

*HSA32* is an HS-inducible gene ubiquitously present in land plants (Charng et al., 2006; Liu et al., 2006). The amino acid sequence of HSA32 shares homology with the archaeal enzyme phosphosulfolactate synthase (ComA), a TIM-barrel protein with a homotrimeric conformation (Wise et al., 2003; Liu et al., 2006). Hence, HSA32 does not belong to the families of HSPs. The molecular function of HSA32 remains unclear, and how it stabilizes HSP101 is unknown. On the other hand, HSP101 was shown to promote the accumulation of HSA32 during the memory phase, indicating a positive feedback loop formed by these two proteins (Wu et al., 2013). The relationship between HSP101 and HSA32 is also present in rice (Lin et al., 2014), suggesting that this positive feedback control is conserved in angiosperms. So far, the precise mechanisms concerning the reciprocal regulation of the two proteins are unclear and remain to be established.

In this study, how HSP101 affects the level of HSA32 was investigated. We observed that different HS temperatures differentially regulated the expression of HSP101 and HSA32, but the interplay between them during the memory phase was not confined to specific temperatures. Degradation of HSA32 was mediated by both autophagy and proteasome in the presence of HSP101. However, HSA32 was degraded much faster in the absence of *HSP101* preferentially via the proteasomal pathway independent of polyubiquitination and the 19S regulatory particle of the 26S proteasome. In response to HS, the newly synthesized HSA32 existed in liquid condensates in the cytosol and was dispersed upon return to non-stress temperatures. To identify HSA32-interacting proteins, we employed biotin ligase TurboID-based proximity labeling, a powerful tool to detect both stable and transient protein interactions within living cells (Branon et al., 2018; Mair et al., 2019). Six potential interacting partners of HSA32 were identified, and HSP101 served as the primary target. The other five potential HSA32-interacting proteins are chaperones and co-chaperones, including three subunits of the eukaryotic cytosolic chaperonin CCT complex and the co-chaperones of HSP70 and HSP90. Intriguingly, these five interactors were also labeled by HSA32-TurboID in the absence of HSP101, the chaperone systems facilitate the folding of the nascent HSA32. We further confirmed that disruption or weakening of the interaction between HSP101 and HSA32 was associated with reduction in HSA32 stability. Moreover, HSP101 affected the biotinylation patterns of HSA32 by TurboID. HSA32 is predicted to be a TIM barrel protein containing intrinsically disordered and aggregation-prone regions. The recombinant HSA32 expressed in *E. coli* was insoluble. These results suggest that HSA32 is prone to aggregate. Altogether, this study provides an example of how plant stress memory can be maintained by a positive feedback loop that, in part, modulates the stability of a heat-inducible aggregation-prone protein by a disaggregase that serves as an AT effector.

## RESULTS

### The protein profiles of HSA32 and HSP101, not their interplay, were affected by priming temperatures

The interplay between HSP101 and HSA32 was revealed during the memory phase after priming at 37°C, at which the accumulation of HSA32 is substantially delayed after priming as compared with other HSPs, including HSP101 (Wu et al., 2013). To determine how priming temperature affects the expression pattern of HSA32 and HSP101, wild type seedlings primed at 32°C, 37°C, and 42°C were analyzed by immune blot. The protein level of HSP101 reached its highest point at early time points during the memory phase after priming at 32°C and 37°C. However, it showed a significant delay in reaching the maximum after acclimation at 42°C. The HSA32 level peaked early at 32°C but delayed substantially above 37°C (Supplemental Figure S1A). To see whether the interplay between the two proteins also exists at different priming temperatures, we examined the protein levels of HSP101 and HSA32 in wild type, *hsp101*, and *hsa32* mutant seedlings primed at 32°C and 42°C. After 32°C heat treatment, HSA32 substantially accumulated at 1 h recovery and sustained about 50% of its maximum level for up to 24 h in wild type. However, in *hsp101*, HSA32 protein started to decline after 1 h recovery and substantially reduced after 4 h (Figure 1A). On the other hand, HSP101 decreased faster in the *hsa32* mutant than in the wild type (Figure 1A). Similar results were observed in the samples primed at 42°C (Supplemental Figure S1B). These results indicate that the interplay between HS101 and HSA32 is not associated with a specific priming temperature. In addition, the interplay occurred in 3-wk-old detached leaves after treatment at 32°C for 1 h (Supplementary Figure S1C), suggesting that this phenomenon not only exists in young seedlings but also in mature plants.

**Figure 1.**
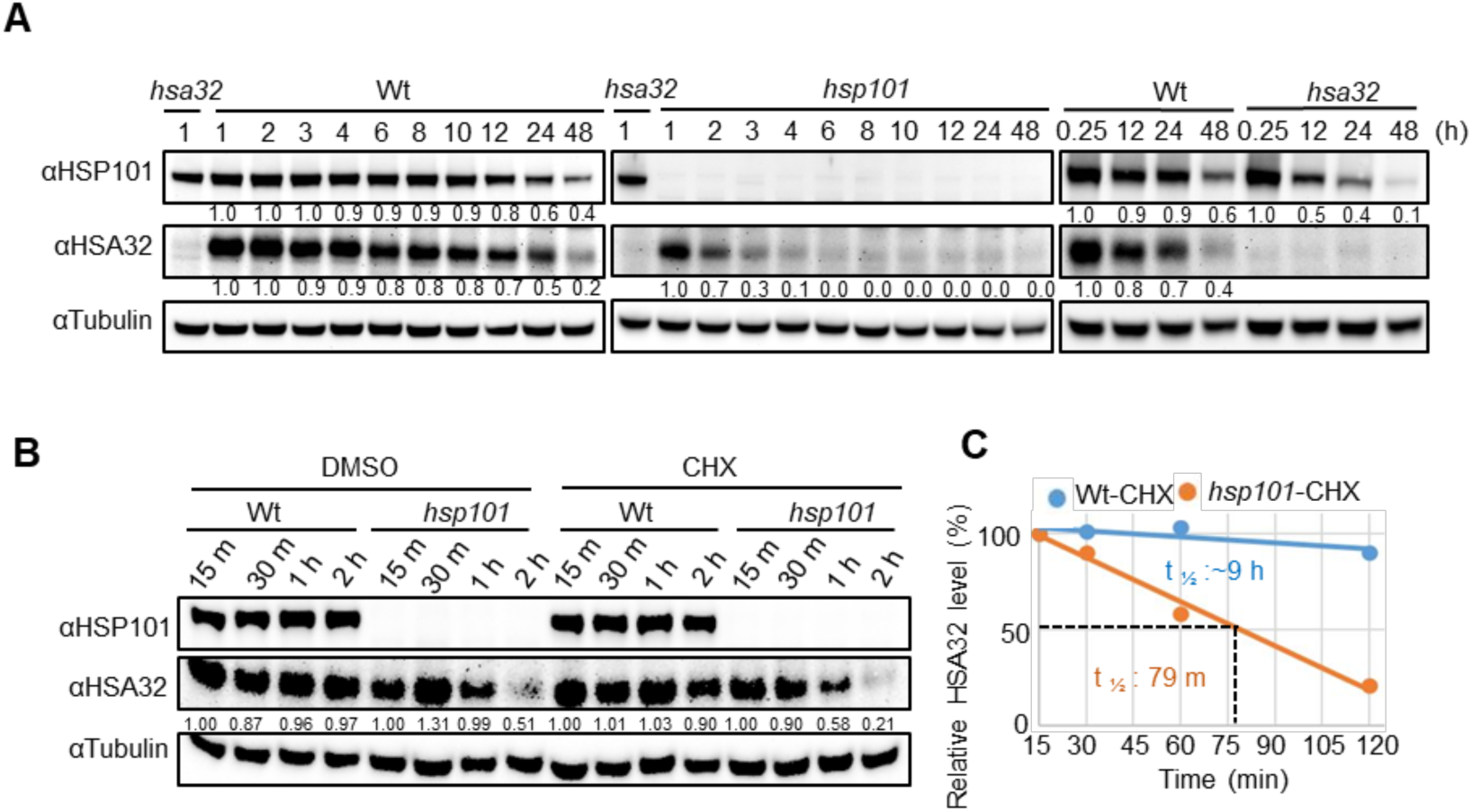
HSP101 protects HSA32 from quick degradation during recovery periods. **A)** The interplay between HSP101 and HSA32 at 32°C in 4 d old seedlings. The protein accumulation pattern of HSA32 was observed in wild type (Wt) and *hsp101* mutant, while the HSP101 protein level was monitored in the wild type and *hsa32*. Each well contained 50 μg proteins, and the blots were probed with antibodies against HSP101, HSA32, and tubulin. ImageJ software was employed to quantify the intensity of the protein bands. **B)** The seedlings (4-d-old) were treated with equal volume (350 µL) of dimethyl sulfoxide (DMSO) and translation inhibitor cycloheximide (CHX) at 15 min of recovery period following heat treatment and collected samples at designated recovery time points. **C)** The half-life of HSA32 was determined using quantified protein band intensity from Figure 1B. The graph’s blue and orange lines represented the *in vivo* decay rate of HSA32 protein in wild type and *hsp101* mutant plants, respectively. Similar results were obtained in three biological repeats.

HSPs, including HSP101 and HSA32, are abundant in the dry seeds. Thus, we examined whether the interplay also occurs at this and subsequent germination stages. We noticed that the levels of these two proteins were reciprocally affected (Supplemental Figure S1D), suggesting that the levels of the two proteins are positively affected by each other at ambient temperature.

Since the settings of heat treatment at 32°C does not substantially delay the accumulation of HSA32 during the memory phase (Supplemental Figure S1A), a majority of the experiments in this study were conducted at this priming temperature to facilitate the analysis of the interplay between HSP101 and HSA32.

### HSP101 prevents HSA32 from rapid degradation mediated by proteasome during memory phase

Since HSA32 could be synthesized shortly after priming at 32°C and disappeared rapidly in *hsp101* (Figure 1A), HSP101 might be involved in the post-translational stabilization of HSA32. To test this hypothesis, we analyzed the decay rate of HSA32 during the memory phase by applying the translation inhibitor, cycloheximide (CHX), to the seedlings after priming. In the presence of CHX, the HSA32 level was slightly affected in the wild type, implying minimal protein degradation. However, a more rapid decline of HSA32 was observed in *hsp101* than in wild type (Figure 1B). The half-life of HSA32 was estimated to be 79 min in *hsp101*, much shorter than that in wild type (Figure 1C), indicating that the degradation of HSA32 was significantly faster in the absence of HSP101.

Two main protein degradation mechanisms in the eukaryotic cells, autophagy and proteasome, were investigated for their role in the rapid degradation of HSA32 in *hsp101*. HSA32 level was monitored in wild type and *hsp101* with blockage of the degradation pathways by introducing an autophagy-defective mutant allele, *atg5*, or by using proteasome inhibitors, MG132 and bortezomib (BTZ). We observed a relatively higher HSA32 protein level in the *atg5* mutant than in the wild type before and after heat treatment, particularly prominent after long recovery (Figure 2A). The protein levels of HSP101 and class I sHSP (sHSP-CI) were examined since these proteins were shown to be degraded via the autophagy pathway (Sedaghatmehr et al., 2019). Indeed, both HSP101 and sHSP-CI were more abundant in *atg5* than in wild type, consistent with the previous report. These results suggest that, in the presence of HSP101, HSA32 can be degraded via the autophagy pathway. To see whether the rapid degradation of HSA32 in *hsp101* might also be mediated by autophagy, we generated *hsp101 atg5* double mutant by crossing the single mutants and compared the decay of HSA32 in *hsp101* and *hsp101 atg5*. No significant difference was observed between *hsp101* and *hsp101 atg5* in HSA32 accumulation (Figure 2B), suggesting that autophagy is not responsible for the rapid degradation of HSA32 in the absence of HSP101.

**Figure 2.**
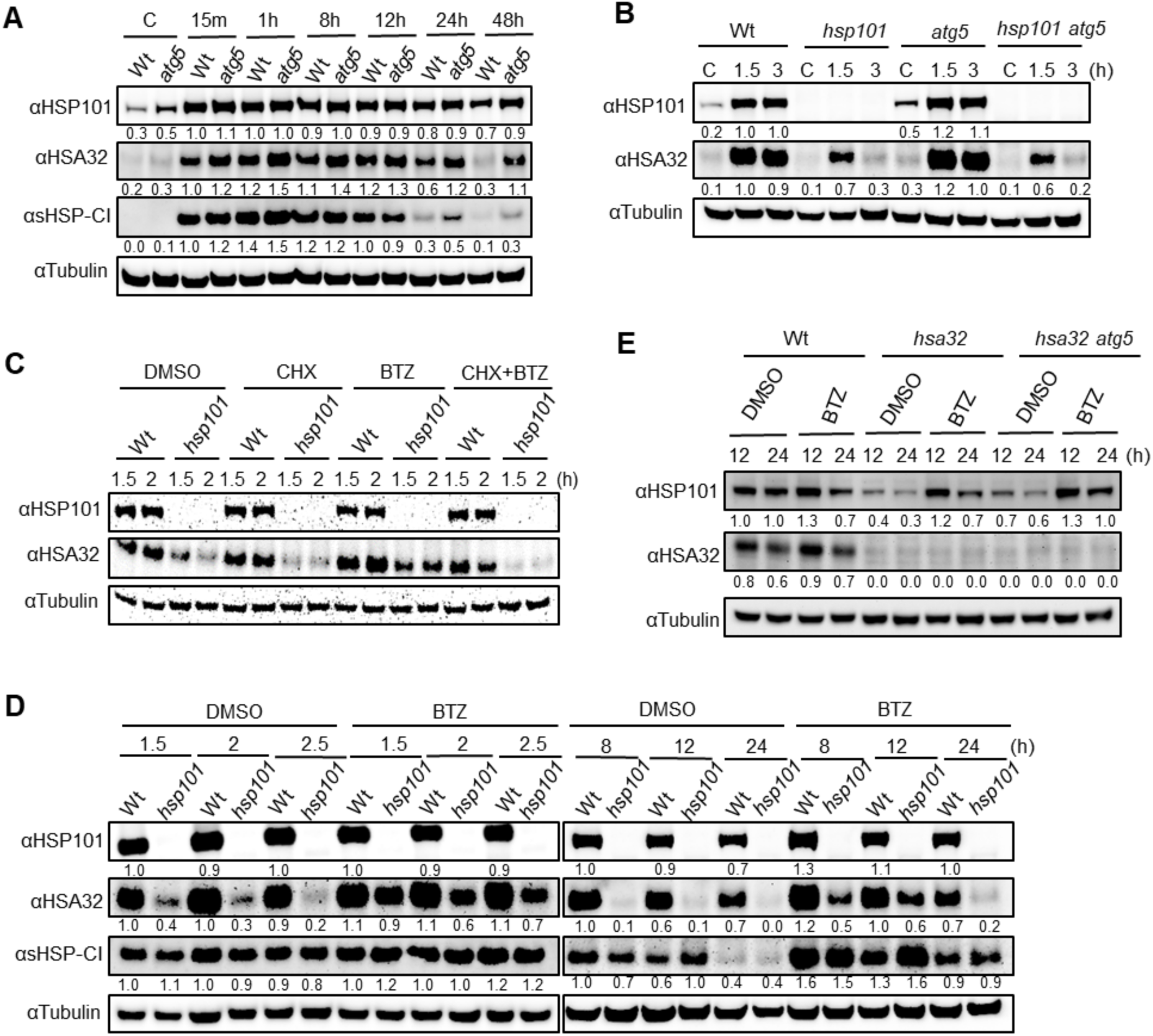
Analysis of HSA32 protein degradation in the plant cell. **A)** To investigate the autophagy-mediated degradation of HSA32, wild type (Wt) and *atg5* seedlings were harvested before and at designated recovery times after heat acclimation at 32°C for 1 h. Antibodies against HSP101, HSA32, sHSP-CI, and tubulin were used to probe the blots. **B)** The protein level of HSA32 was further observed in wild type, *hsp101*, *atg5*, and double mutant *hsp101 atg5*. **C)** The wild type and *hsp101* mutant seedlings were treated with 0.1% DMSO (control), the translation inhibitor (50 µM CHX), the proteasome inhibitor (50 µM BTZ), and a combination of both CHX and BTZ in the Petri dish following heat acclimation treatment and collected samples at 1.5 and 2 h recovery times and analyzed by immunoblotting. **D)** The DMSO and BTZ treated wild type and *hsp101* seedlings were allowed to recover for a wide range of designated times and collected samples for immunoblotting to validate the proteasome-mediated degradation of HSA32. Three independent biological replicates yielded consistent results. **E)** The degradation pathway of HSP101 was examined in the presence and absence of BTZ using *hsa32* single and *hsa32 atg5* double mutants and analyzed by immunoblotting with anti-HSP101 and anti-HSA32 antibodies. ImageJ software was employed to quantify the intensity of the protein bands.

Then, the role of proteasome was assessed. Our results show that BTZ, but not MG132, could substantially block the degradation of HSA32 in *hsp101* (Supplemental Figure S2A). Intriguingly, BTZ could not promote the accumulation of HSA32 in *hsp101* in the presence of CHX (Figure 2C), suggesting that the nascent HSA32 protein was susceptible to proteasome without HSP101. Afterward, we validated the effect of BTZ without CHX on the degradation of HSA32 during a wide range of recovery periods after priming. As observed, BTZ treatment substantially increased the HSA32 level in *hsp101* as compared to the dimethyl sulfoxide (DMSO) solvent control during the memory phase, particularly pronounced within 1.5 to 2.5 h (Figure 2D). This finding implies that the rapid degradation of HSA32 in *hsp101* occurs through the proteasome system.

Interestingly, BTZ noticeably suppressed the degradation of HSP101 and sHSP-CI at the late memory phase (8-24 h, Figure 2D), suggesting that both HSP101 and sHSP-CI are degraded through the proteasome system. However, proteasome-mediated degradation of sHSP-CI was not affected by HSP101 as the protein level shows no significant difference between wild type and *hsp101*, indicating the selectivity of HSP101’s effect on HSA32 stabilization. We further strengthen this result by noticing that both *hsp101* and *hsp101 atg5* exhibited approximately similar trends of HSA32 protein accumulation in the BTZ-treated samples as compared to the DMSO control (Supplemental Figure S2B), implying that the proteasome pathway plays a dominant role for the fast degradation of HSA32 in the absence of HSP101.

We noticed that the HSA32 protein level in *hsp101* could not be restored to the wild-type level when BTZ was applied after the heat treatment (Figure 2, C and D), which might be due to the timing of the application of the inhibitor. We applied BTZ to the seedlings one hour or immediately before the heat acclimation treatment to verify this hypothesis. Similar HSA32 levels were observed in wild type and *hsp101* with BTZ applied before but not after the heat treatment (Supplemental Figure S2C), suggesting that the nascent HSA32 is quickly degraded via the proteasome pathway in the absence of HSP101.

Since HSP101 could be degraded by autophagy and proteasome in wild type (Figure 2, A and D), we examined whether the faster degradation of HSP101 in *hsa32* might occur via the two pathways. Our results showed that the HSP101 level was slightly higher in *hsa32 atg5* than in *hsa32* but lower than in the wild type after 12 and 24 h of recovery (Figure 2E), suggesting that HSA32 prevents HSP101 from autophagy-mediated degradation. BTZ treatment further reduced the degradation of HSP101 in *hsa32 atg5* and *hsa32* (Figure 2E), indicating that, in addition to autophagy, the faster degradation of HSP101 in *hsa32* also involved the proteasome pathway.

### The role of polyubiquitination and 26S proteasome in HSA32 degradation

To examine whether HSA32 is polyubiquitinated for proteasome-mediated degradation, we performed the Tandem Ubiquitin-Binding Entities (TUBE) assay on transgenic lines HA-tagged HSA32. The HA-HSA32 successfully complemented *hsa32* by rescuing the LAT defect (Supplemental Figure S2D). To assess the stability of HA-HSA32 without HSP101, we crossed the transgenic lines with *hsp101 hsa32* mutant. We observed that HA-HSA32 was less stable in the absence of HSP101 and was stabilized by BTZ (Supplemental Figure S2E), suggesting that it is degraded by the proteasome like the endogenous HSA32. Lys48-polyubiquin-bound proteins were purified using the TUBE resin in the presence and absence of BTZ. Polyubiquitinated proteins accumulated more when BTZ was applied, particularly in the absence of HSP101 (Supplemental Figure S2F). Unfortunately, this experiment could not detect any significant high-molecular weight signals representing polyubiquitinated HA-HSA32, suggesting that HA-HSA32 might not be polyubiquitinated (Supplemental Figure S2F). This result was further verified using PYR-41, an inhibitor of the ubiquitin-activating enzyme E1, to block protein polyubiquitination and 26S proteasome action. We found that PYR-41 could not suppress the fast degradation of HSA32 in *hsp101* (Supplemental Figure S2G). Instead, it slightly promoted HSA32 degradation. As a positive control, the degradation of NPR1-GFP, which is mediated by the ubiquitin–proteasome system (Skelly et al., 2019), was suppressed by PYR-41 (Supplemental Figure S2H). Taken together, these results suggest that HSA32 is degraded by proteasome without being polyubiquitinated.

To confirm the involvement of 26S proteasome in HSA32 degradation, we employed a genetic approach using the mutants of *RPN10*, a subunit of the 19S regulatory particle of the 26S proteasome, that results in reduced 26S proteasome activity. RPN10 serves as the primary ubiquitin chain receptor, facilitating the degradation of the substrate by 26S proteasome (Lin et al., 2011; Martinez-Fonts et al., 2020). We hypothesized that if HSA32 underwent degradation via the 26S proteasome, we would observe a higher accumulation of HSA32 protein in the *RPN10* mutants. Consistent with the previous reports (Kurepa et al., 2008; Lin et al., 2011), the *RPN10* mutants exhibited increased levels of polyubiquitinated proteins and 20S proteasome as compared with the wild types (Col-0 and C-24, Supplemental Figure S2I). However, contrary to our hypothesis, a relatively smaller amount of HSA32 protein was observed in the *RPN10* mutants compared to the wild types after priming at 32°C. HSP101 levels were not affected in the mutants. These results suggest that the 19S regulatory particle of the 26S proteasome is not involved in the degradation of HSA32, consistent with the observations that HSA32 degradation was not associated with polyubiquitination.

### HSA32 is mainly located in the cytosol and transiently associates with heat-induced condensates

HSP101 is known to be located in the cytoplasm (Lee et al., 2007; McLoughlin et al., 2019). However, the localization of HSA32 remained unclear and is predicted to be located in the nucleus (The Arabidopsis Information Resource, www.arabidopsis.org/servlets/TairObject?id=130253&type=locus, Oct 4, 2023). To verify this prediction, we performed subcellular fractionation analysis by differential ultracentrifugation of the cell lysate of wild type seedlings harvested after heat acclimation. Our result showed that HSA32 was barely detectable in the nucleus fraction and mainly in the cytosolic fraction (Figure 3A). A similar distribution pattern was observed for HSP101, suggesting that both are mainly in the cytosol.

**Figure 3.**
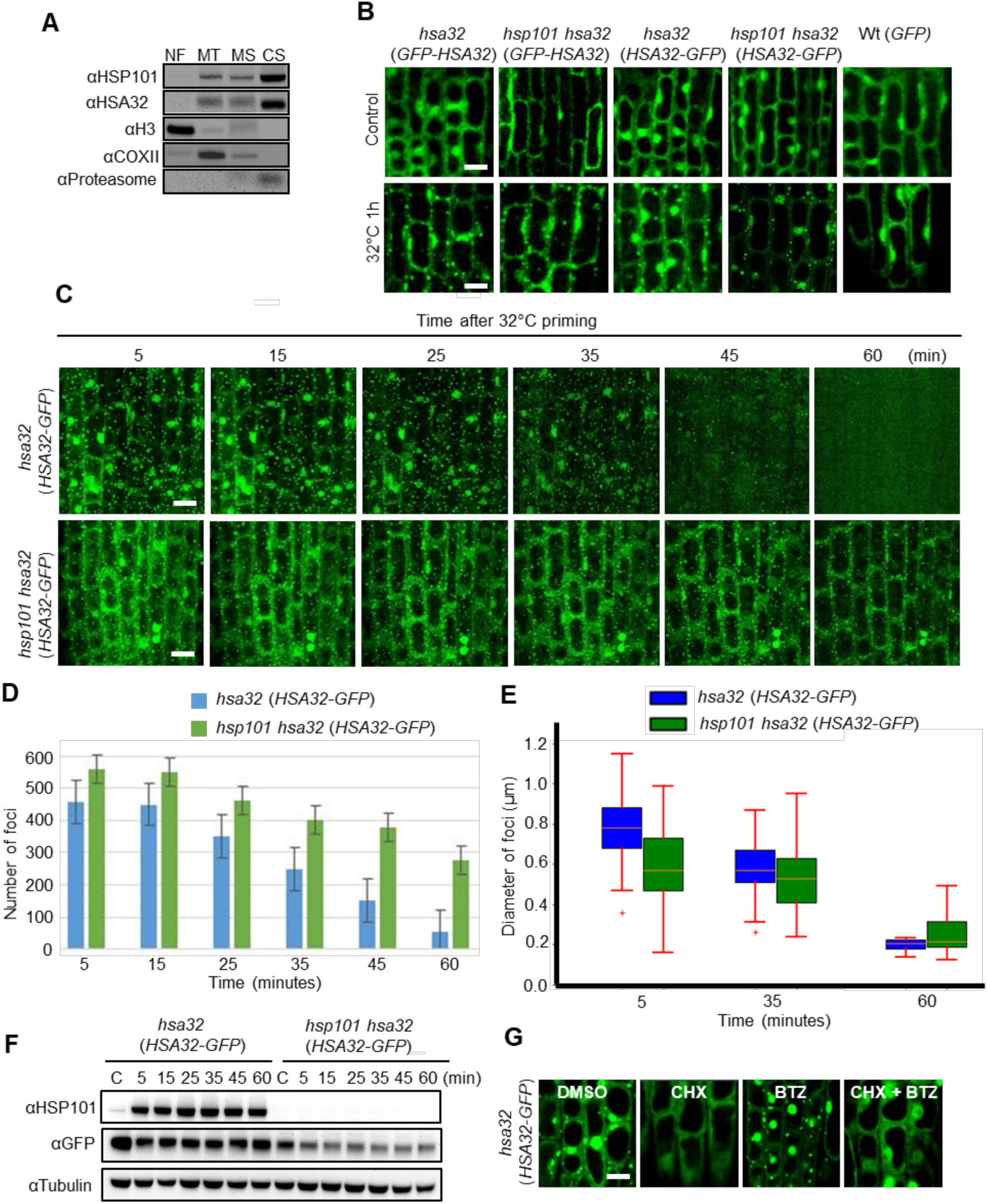
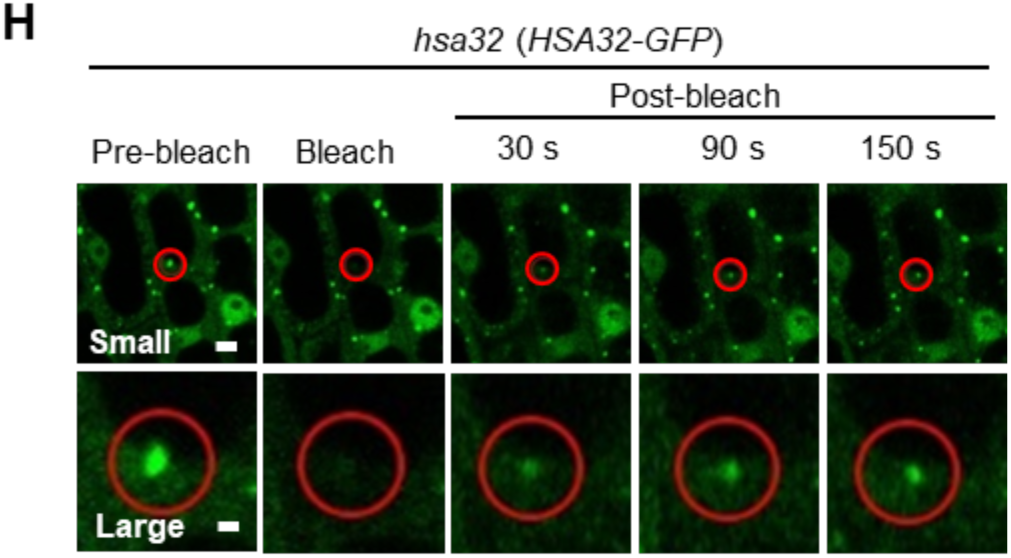
Subcellular localization and foci formation of HSA32. **A)** Subcellular compartments of heat-acclimated wild type seedlings were fractionated using differential ultra-centrifugation of the cell lysates. Four distinct fractions were collected and analyzed: nuclear fraction (NF), mitochondrial fraction (MF), microsomal fraction (MS), and cytosolic soluble fraction (CS). Anti histone3 (H3), COXII, and proteasome antibodies were used as markers for nuclear, mitochondrial, and cytosolic soluble fractions, respectively. **B)** Confocal microscopy images of stably expressing *p35S*:*HSA32-GFP* in *hsa32* and *hsp101 hsa32* background and *p35S*:*GFP* in wild-type background. The images of root cells were taken under the control and after heat stress at 32°C for 1 h. Scale bars, 10 μm. **C)** Time-lapse imaging of HSA32-GFP foci assembly and disassembly with or without HSP101 under indicated recovery times following heat stress at 32°C for 1 h. Scale bars, 5 μm. The number of foci was quantified by ImageJ. **D)** The graph depicted the number of foci observed in *hsa32* (*HSA32-GFP*) and *hsp101 hsa32* (*HSA32-GFP*), represented by the blue and green colors, respectively, during designated recovery intervals. Notably, the number of foci disappeared faster in the presence of HSP101. **E)** The box and whisker plots represent the diameter of HSA32 foci (n =100 foci at 5 min and 35 min each, and n = 35 foci at 60 min) from multiple cells. The central line represents the median value, the line inside the box indicates the average value, the box encompasses the upper and lower quartiles, and the error bars show the maximum and minimum size distributions. **F)** The protein level of HSA32 was examined at indicated recovery periods by immunoblotting, and HSA32-GFP protein gradually decreased in the absence of HSP101. **G)** Seedlings were treated with 0.1% DMSO, 50 µM CHX, 50 µM BTZ, and a combination of CHX and BTZ before 1 h of heat stress and observed under a confocal microscope immediately after heat stress at 32°C. Scale bars, 10 μm. **H)** FRAP (<u>F</u>luorescence <u>r</u>ecovery <u>a</u>fter <u>p</u>hotobleaching) analysis of *hsa32* (*HSA32-GFP*) foci upon heat stress. The red circle represented the targeted foci for bleaching, and post-bleaching was monitored at the timeframes mentioned after bleaching. The upper and lower panels represented the results of FRAP analysis with small and large images, respectively. Scale bars, 2 μm. All confocal images are representative of three independent biological replicates.

To monitor the location of HSA32 *in vivo*, we employed the transgenic plants expressing GFP-tagged HSA32 in *hsa32* under the control of the CaMV35S promoter. The GFP tag was fused to the N- or C-terminus of HSA32 to yield GFP-HSA32 and HSA32-GFP, respectively. The recombinant proteins rescued the LAT defect of *hsa32* (Supplemental Figure S3A), suggesting that the fusion proteins are functional. To assess the effect of HSP101 on the location of HSA32, we introduced the transgenes to the *hsp101 hsa32* background by genetic crossing of the transgenic lines and the double mutant. In the absence of HSP101, the GFP-HSA32/HSA32-GFP fusion proteins were accumulated to lower levels as compared with that in the presence of HSP101 (Supplemental Figure S3B), suggesting that HSP101 stabilized the fusion proteins like the native protein. Subcellular fractionation analysis of the cell lysate of the transgenic plants (in *hsa32* background) showed that GFP-HSA32 was also mainly located in the cytosol with a small amount in the nucleus fraction (Supplemental Figure S3C).

Under non-stress conditions, GFP-HSA32 and HSA32-GFP fluorescent signals were diffusely distributed in the cytosol and nucleus with or without HSP101, similar to the GFP control (Figure 3B). Intriguingly, following heat treatment at 32°C for 1 h, condensed fluorescent foci were observed in the cytosol of the fusion protein expressing cells, irrespective of HSP101 (Figure 3B), suggesting that HSP101 is dispensable for the foci formation. GFP alone did not form such foci in the control line, suggesting that HSA32 is responsible for the foci formation upon heat treatment.

Time-lapse imaging showed that the HSA32-GFP foci appeared after 30 min of heat treatment (Supplemental Figure S3, D and E) and gradually disappeared during the memory phase, with the one with HSP101 present seemingly disappearing faster than its absence (Figure 3C). The foci number seemed higher in the absence than in the presence of HSP101 immediately after heat stress (Figure 3D). The size of HSA32 foci appeared significantly more prominent in the presence than in the absence of HSP101 (Figure 3E). Immunoblots showed that the amount of HSA32-GFP remained steady within 60 min, and this protein declined in the absence of HSP101 (Figure 3F). The results suggest that the variations in foci number were not correlated with HSA32-GFP protein abundance, and similar results were observed in Supplemental Figure S3, D and F. To see whether the foci were associated with newly translated or preexisting HSA32-GFP, we examined the foci formation by applying individually and in combination with CHX and BTZ. Notably, HSA32-GFP foci formation was entirely abolished by CHX or by CHX plus BTZ; BTZ treatment did not prevent foci formation (Figure 3G). This result suggests that HSA32 foci are associated with the newly synthesized HSA32.

Fluorescence recovery after photobleaching (FRAP) assay was conducted to determine the nature of the HSA32-GFP foci. We observed that the condensed HSA32-GFP signals gradually recovered after bleaching within the bleached areas (Figure 3H), suggesting that the foci are liquid condensates formed through liquid-liquid phase separation (LLPS). The foci did not fuse but gradually vanished during the memory phase (Supplemental Figure S3G). Next, we explored whether the HSA32-GFP foci are, in essence, stress granules (SGs) induced by heat and formed through LLPS (Wheeler et al., 2016; Ruiz-Solaní et al., 2023). The SG marker, PAB2-RFP, formed foci at 37°C but not 32°C (Supplemental Figure S3H). However, HSA32 foci appeared at 32°C and decreased in number at 37°C (Supplemental Figure S3H), suggesting that they are not the same type of liquid condensates.

### Identification of potential HSA32-interacting proteins by proximity labeling and LC-MS/MS analysis

To understand how HSP101 specifically stabilizes HSA32, we examined whether this action requires direct interaction of the two proteins. We employed various methods like yeast-two-hybrid (Y2H), bimolecular fluorescence complementation (BiFC), and co-immunoprecipitation (Co-IP) assays. Attempts in Y2H and BiFC assays failed as the constructs with HSA32 fusion led to false positive results. We used transgenic plants expressing HSA32-GFP for the Co-IP assay to see if HSP101 can co-immunoprecipitate by GFP antibodies. However, we could not obtain positive results either (Supplemental Figure S4A).

The failure in the Co-IP assay may be due to transient interaction between HSA32 and HSP101. Thus, TurboID-based proximity labeling was employed. We generated recombinant DNAs that encode HA-tagged TurboID (an improved BirA* biotin ligase variant) fused to HSA32 at either the N-terminus (HA-TurboID-HSA32) or C-terminus (HSA32-HA-TurboID). The transgenic plants expressing HA-TurboID, shown to be located in the cytosol and nucleus (May et al., 2020), were generated as the reference control. These recombinant genes were all under the control of the native promoter of *HSA32*. The transgenes were delivered into *hsa32* to generate stable transgenic lines designated as *hsa32* (*HA-TurboID-HSA32*), *hsa32* (*HSA32-HA-TurboID*), and *hsa32* (*HA-TurboID*). The LAT defect of *hsa32* was rescued to some extent in all independent lines of *hsa32* (*HA-TurboID-HSA32*) and *hsa32* (*HSA32-HA-TurboID*) but not in *hsa32* (*HA-TurboID*) (Supplemental Figure S4B). The expression of the recombinant proteins was confirmed by immunoblot analysis of the transgenic lines. All the TurboID-containing proteins were expressed to similar levels and were slightly increased after heat treatment (Supplemental Figure S4C). Next, the transgenic lines were crossed with *hsp101 hsa32* to observe the stability of the fusion proteins in the absence of HSP101. HA-TurboID-HSA32 and HSA32-HA-TurboID became unstable without HSP101, while HA-TurboID was unaffected (Supplemental Figure S4D). These results suggest that the HSA32 fusion proteins are functional and can form a positive feedback loop with HSP101 in maintaining AT.

First, to identify the biotinylated proteins, the transgenic lines in the *hsa32* background were incubated with different biotin concentrations and durations (Figure 4A). A distinct biotinylated protein band near 100 kDa appeared in the sample expressing TurboID fused to the C-terminus of HSA32. However, HA-TurboID-HSA32 and the TurboID control did not exhibit such a band (Figure 4A). Furthermore, the 100 kDa band was missing in the line expressing HSA32-HA-TurboID in the *hsp101 hsa32* background, in which the endogenous HSP101 and HSA32 were eliminated through crossing (Figure 4B). This genetic evidence suggests that this band was biotinylated HSP101.

**Figure 4.**
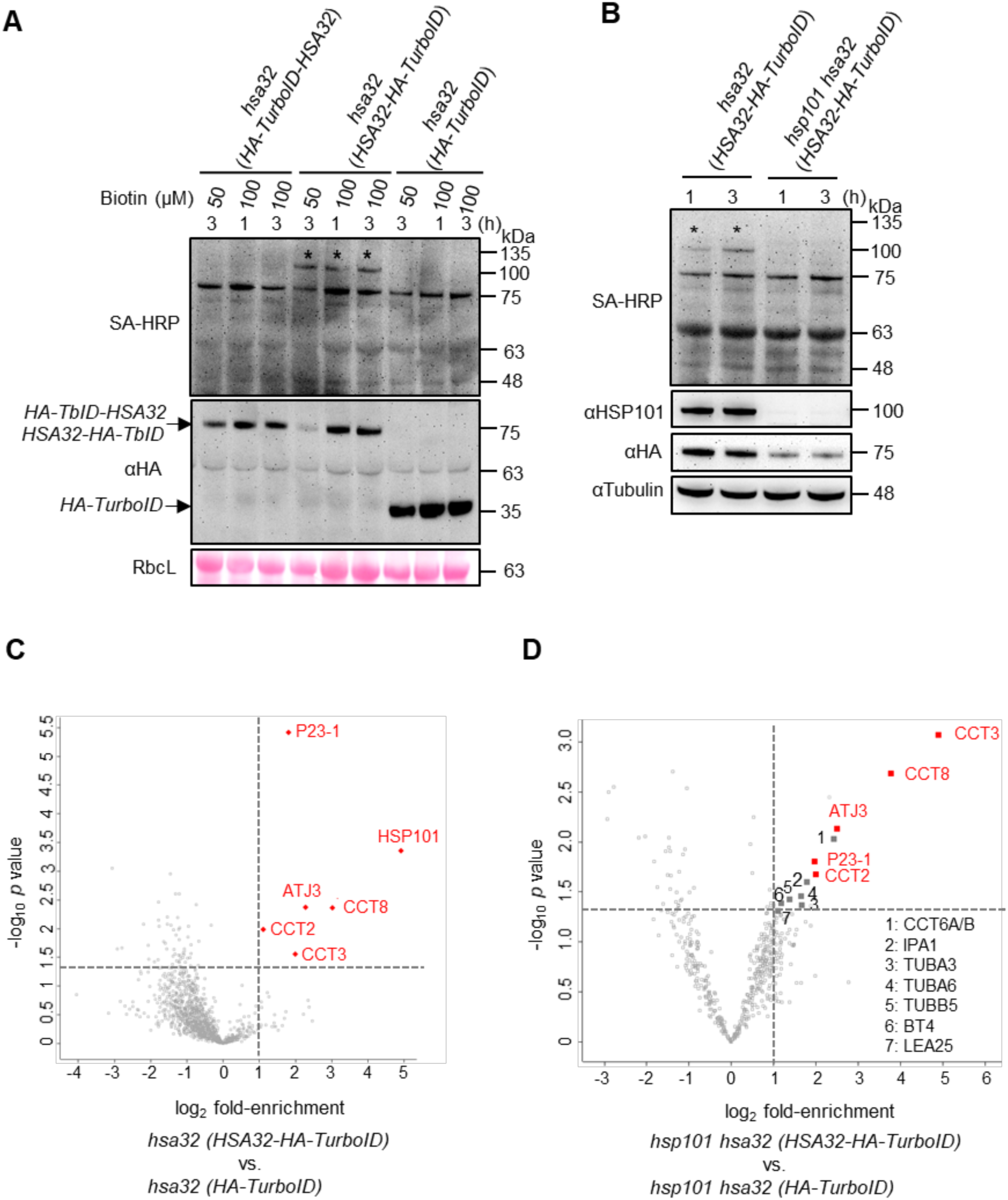
Identification of HSA32 interacting proteins by TurboID-based proximity labeling. **A)** HA-TurboID fused with HSA32 showed biotinylation after being treated with exogenous biotin at 50 and 100 µM concentrations for labeling times of 1 and 3 h of 4-d-old seedlings. Streptavidin-horseradish peroxidase conjugate (SA-HRP) was used to detect the biotinylated proteins. In immunoblotting analysis, an anti-HA antibody was employed to identify HA-TurboID fused HSA32 or HA-TurboID. RbcL was used as a loading control. The molecular weights (kDa) were indicated on the right, and the size of TurboID fusion proteins was shown on the left side with arrows. Biotinylated bands near 100 kDa were marked with asterisks. The experiment was repeated three times with similar results. **B)** HSA32-HA-TurboID in *hsa32* single mutant and *hsp101 hsa32* double mutant’s background were used to check the biotinylated bands after applying 100 µM biotin exogenously on the seedlings. The blots were probed with SA-HRP to examine the biotinylated proteins, and an antibody against HSP101 was used to confirm the abundance of HSP101 protein. The volcano plot visualized significantly enriched proteins in HSA32-HA-TurboID in *hsa32* background **(C)** and *hsp101 hsa32* background **(D)** with the *p*-value less than 0.05 and a greater than 2-fold enrichment based on comparisons between HSA32-HA-TurboID and HA-TurboID. **D)** Proteins depicted in red colored represent interactors that were commonly identified in both the presence and absence of HSP101. Conversely, proteins colored in grey represent those that were newly enriched and exclusively identified only in the samples without HSP101. All data were collected from three independent biological replicates.

Next, the total biotinylated proteins were purified from the extracts of heat-acclimated *hsa32* (*HSA32-HA-TurboID*) and *hsa32* (*HA-TurboID*) using streptavidin-conjugated resin after confirming equal expression of the transgenes and loading of total proteins (Supplemental Figure S4E). The purified biotinylated protein samples were then trypsin-digested and subjected to LC-MS/MS analysis. Venn diagrams showed a high degree of overlap (> 93%) of the hits between the HSA32-HA-TurboID and HA-TurboID samples in each replicate (Supplemental Figure S4F). However, less than 40% of the hits were shared among three biological replicates (Supplemental Figure S4G), suggesting a relatively high degree of non-reproducible noise from experiments performed on different dates. GO cellular component of the identified proteins indicated that nuclear and cytosolic proteins were the most enriched in HSA32-HA-TurboID and HA-TurboID samples (Supplemental Figure S4H). The similar distribution of GO cellular components indicates that HA-TurboID samples serve as reliable references for those of HSA32-HA-TurboID without generating substantial biases due to differences in compartmentalization.

To identify potential HSA32 interacting proteins, we implemented two filtering criteria: the abundance ratio of the identified protein in *hsa32* (*HSA32-HA-TurboID*) versus *hsa32* (*HA-TurboID*) ≥ 2 and the p-value < 0.05 in all three biological replicates. We observed that HSA32 peptides were identified in *hsa32* (*HSA32-HA-TurboID*) but not in *hsa32* (*HA-TurboID*), suggesting inter- or intra-molecular labeling of the HSA32-HA-TurboID protein (Supplemental Table S1). Volcano plot analysis showed, in addition to HSA32 itself, six potential interactors with significantly increased abundance due to the HSA32 fusion (Figure 4C). Among them, HSP101 exhibited the highest abundance ratio and was identified with the highest number of unique peptides and PSM value (Supplemental Table S2). In addition to HSP101, the other five potential interactors are also chaperones or co-chaperones, including three subunits of the chaperonin-containing TCP1 (CCT) complex (CCT2, CCT3 and CCT8) and two cytosolic co-chaperones, P23-1 and AtJ3 (Supplemental Table S2).

Furthermore, the potential interactors of HSA32 in the absence of HSP101 were identified from the extracts of the transgenic lines expressing HSA32-HA-TurboID and HA-TurboID in the *hsp101 hsa32* background. Like the case of the transgenic lines in the *hsa32* background, the numbers of hits were similar between the *hsp101 hsa32* (*HSA32-HA-TurboID*) and *hsp101 hsa32* (*HA-TurboID*) samples with a high degree of overlap (> 94%) in all three biological replicates (Supplemental Figure S4I). More than 95% of the hits were shared among three biological replicates (Supplemental Figure S4J). GO cellular component analysis demonstrated similar patterns for *hsp101 hsa32* (*HSA32-HA-TurboID*) and *hsp101 hsa32* (*HA-TurboID*) (Supplemental Figure S4K). Notably, 465 proteins (43%) were uniquely identified in the presence of HSP101, while 229 proteins (21%) were unique to samples without HSP101 and a substantial overlap of 389 identified proteins (36%) (Supplemental Figure S4L).

To identify potential HSA32 interactors without HSP101, we applied the same filtering criteria mentioned in Figure 4C. HSA32 peptides were identified in *hsa32 hsp101* (*HSA32-HA-TurboID*) but not in *hsa32 hsp101* (*HA-TurboID*) (Supplemental Table S3). Volcano plot analysis showed, in addition to HSA32 itself, twelve potential HSA32 interactors that fulfilled the criteria (Figure 4D). Interestingly, all of the potential interactors identified in *hsa32* (*HSA32-HA-TurboID*) except HSP101 were also significantly enriched in *hsp101 hsa32* (*HSA32-HA-TurboID*), suggesting that these proteins interact with HSA32 in an HSP101-independent manner. CCT6A/B, IPA1, TUBA3, TUBA6, TUBB5, BT4, and LEA25 were solely enriched when HSP101 was absent (Supplemental Table S4).

HSA32 was found to be one of the proteins enriched by the streptavidin-conjugated resin in the presence and absence of HSP101 (Supplemental Table S2 and 4), suggesting that HSA32-HA-TurboID can biotinylate the HSA32 portion intermolecularly or intramolecularly.

### HA tagging compromised HSP101 function and its biotinylation by HSA32-HA-TurboID

Previous studies have shown that the N-terminal domain of ClpB and HSP104 serves as the substrate-binding site, and a tag fused to the N-terminus of ClpB was shown to interfere with its function (Chow et al., 2005; Rosenzweig et al., 2015). HSP101 with a GFP tag at its C-terminus was shown to complement *hsp101* in SAT (McLoughlin et al., 2019), suggesting that the C-terminal tag would not affect HSP101’s function. We confirmed that the expression of HSP101 with an N-terminal HA tag (HA-HSP101) could not rescue the mutant in both SAT and LAT assays (Figure 5A), suggesting that HA-HSP101 lost the chaperone function like the N-terminally tagged ClpB. However, expression of HSP101 with a C-terminal HA tag (HSP101-HA) could rescue the SAT but not LAT defect of *hsp101* (Figure 5A), suggesting that the C-terminal tag did not disrupt the chaperone function of HSP101 but affected the maintenance of AT. Immunoblot analysis showed that both HA-HSP101 and HSP101-HA failed to maintain the levels of HSA32 and, subsequently, their abundance after a long memory phase (Figure 5B and C), suggesting that the HA-tagged HSP101s do not stabilize HSA32 as effectively as the wild type HSP101.

**Figure 5.**
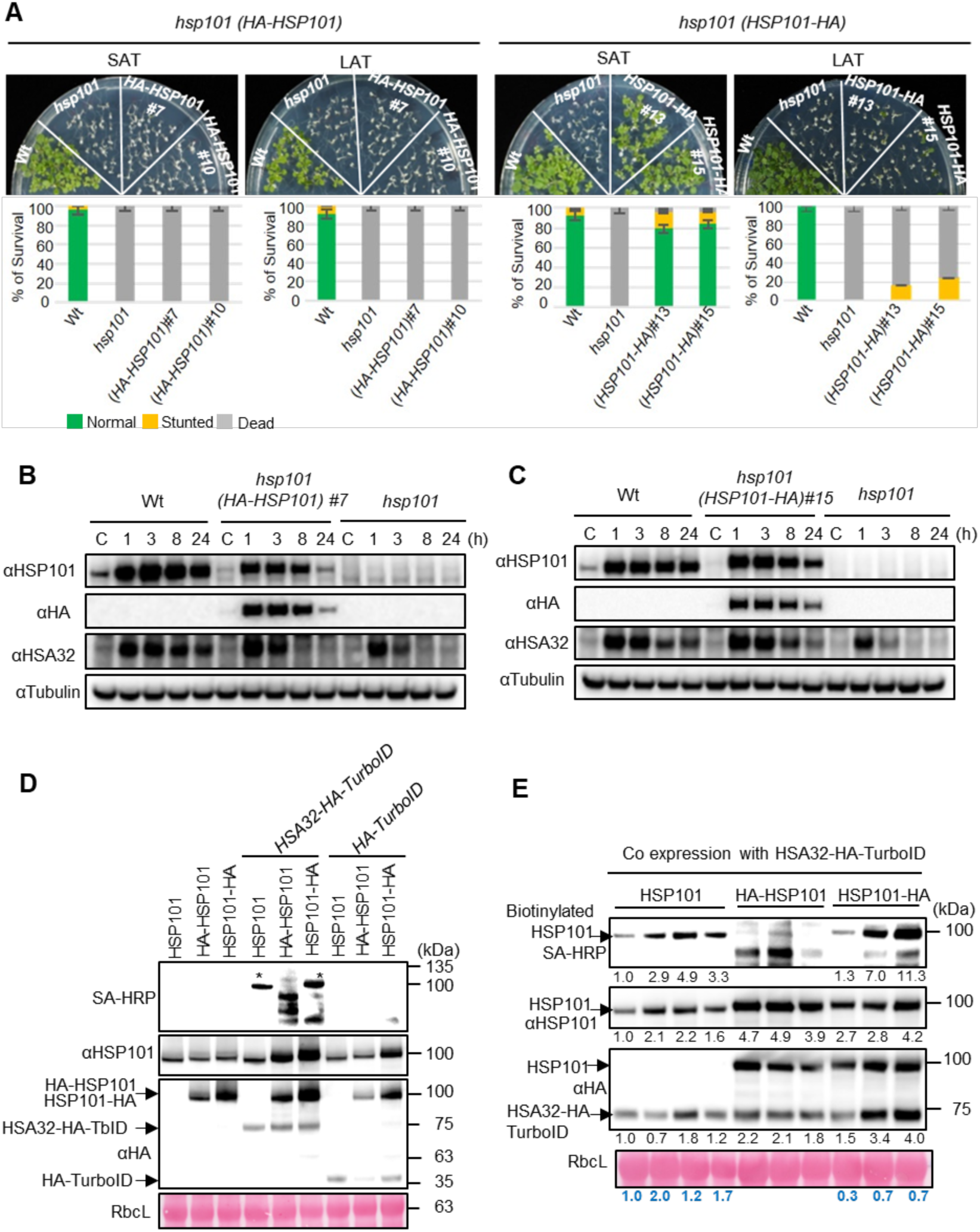
Interaction between HSP101 and HSA32 is associated with HAM. **A)** The results of the SAT and LAT assays for the complementation lines expressing HSP101 variants with HA tag fused to the N-and C-terminus (HA-HSP101) and (HSP101-HA) in *hsp101* background. Immunoblot analyses of HA-HSP101 (**B**), HSP101-HA (**C**), and HSA32 levels followed HS at 32°C for 1 h with indicated recovery periods. The protein levels of HSP101 and HSA32 were monitored using antibodies against HSP101, HA tag, and HSA32. **D)** HSA32-HA-TurboID and HA-TurboID were co-expressed with Arabidopsis HSP101 alone or with HA-HSP101 or HSP101-HA in *N. benthamiana* leaves. Co-infiltration of agrobacteria was conducted in the leaves of three-week-old plants, followed by biotin (200 μM) infiltration into the same leaf sectors one day after agroinfiltration. The infiltrated leaves were collected for protein extraction one day after biotin infiltration. SA-HRP and antibodies (against HSP101 and HA tag) were used to probe the blots. The asterisks marked for the biotinylated proteins were approximately 100 kDa in size. **E)** The interaction efficiency between HSP101 and HSA32-HA-TurboID was compared to that between HSP101-HA and HSA32-HA-TurboID. The expression levels of HSP101, HA-HSP101, HSP101-HA, and HSA32-HA-TurboID were modulated by the concentration of agrobacteria harboring the transgenes. The specific combinations of agrobacteria concentrations are detailed in Supplemental Table S5. The relative intensity of biotinylated and non-biotinylated HSP101 as well as HSA32-HA-TurboID protein bands were quantified by ImageJ and displayed below each corresponding gel image. The efficiency of the biotinylation reaction was calculated using the following equation: biotinylated HSP101 signal/(HSP101 protein signal x HSA32-HA-TurboID protein signal).

Next, we employed agro-infiltration to transiently co-express HSA32-HA-TurboID and the HSP101 variants in the tobacco leaves to examine their interaction. HSA32-HA-TurboID protein levels were lower when expressed alone in the tobacco leaves than when co-expressed with wild type HSP101, suggesting that HSP101 stabilizes HSA32-HA-TurboID in tobacco cells. A biotinylated band around 100 kDa appeared when HSP101 was co-expressed with HSA32-HA-TurboID, not with HA-TurboID (Supplemental Figure S5), which is similar to the results of the stable transgenic plants (Figure 4A). However, the biotinylation of HA-HSP101 was substantially reduced, while that of HSP101-HA was unaffected (Figure 5D). As a control, the HSP101 variants co-expressed with HA-TurboID did not result in the corresponding biotinylated bands (Figure 5D). These results suggest that the labeling of HSP101 by HSA32-HA-TurboID was disrupted by HA-tag fused to the N-terminus of HSP101. Of note, additional biotinylated protein bands of smaller sizes were observed in the sample expressing HA-HSP101. To further compare the labeling efficiency of native HSP101 and HSP101-HA by HSA32-HA-TurboID, we manipulated the expression of the proteins by adjusting the dose of co-injected agrobacterium solutions (Supplemental Table S5). Since the level of the biotinylated HSP101/HSP101-HA was positively correlated with the amount of HSP101/HSP101-HA and HSA32-HA-TurboID, we estimated the efficiency of the biotinylation reaction by the ratio of biotinylated HSP101 signal/(HSP101 protein signal x HSA32-HA-TurboID protein signal). Our data showed that HSP101 was more efficiently labeled (the ratio was between 1.0 and 2.0) than HSP101-HA (the ratio was between 0.3 and 0.7), suggesting that the HA-tag fused to the C-terminus of HSP101 interferes its biotinylation. Again, HSA32-HA-TurboID labeled HA-HSP101 could not label HA-HSP101 (Figure 5E). These results suggest the necessity of protein-protein interaction for their biological function in maintaining AT.

### Differential biotinylation of HSA32 by TurboID in the presence and absence of HSP101

Since TurboID preferentially biotinylates surface-exposed lysine residues (Shioya et al., 2022), protein conformational status may affect the biotinylation pattern. Thus, we analyzed and compared the biotinylation sites of HSA32 in the presence and absence of HSP101 using the LC-MS/MS data generated in Figure 4. Our results show that K8, K67, K138, K145, K154, K220, and K229 were biotinylated among 18 lysine residues of HSA32 (Figure 6) and (Supplemental Table S6). Notably, biotinylation was exclusively detected at K8, K138, and K145 only when HSP101 was present (Figure 6). In contrast, the biotinylation of K67, K154, K220, and K229 was irrespective of the presence or absence of HSP101 (Figure 6). As expected, all the identified biotinylated lysine residues (except K145) were located on the surface of the 3D structure of the predicted monomeric HSA32 (Supplemental Figures S6A and S6B). These results suggest that HSP101 affects the accessibility of K8, K138, and K145 on HSA32, and in the presence of HSP101, these residues are susceptible to biotinylation inter- or intramolecularly by the TurboID fused to HSA32.

**Figure 6.**
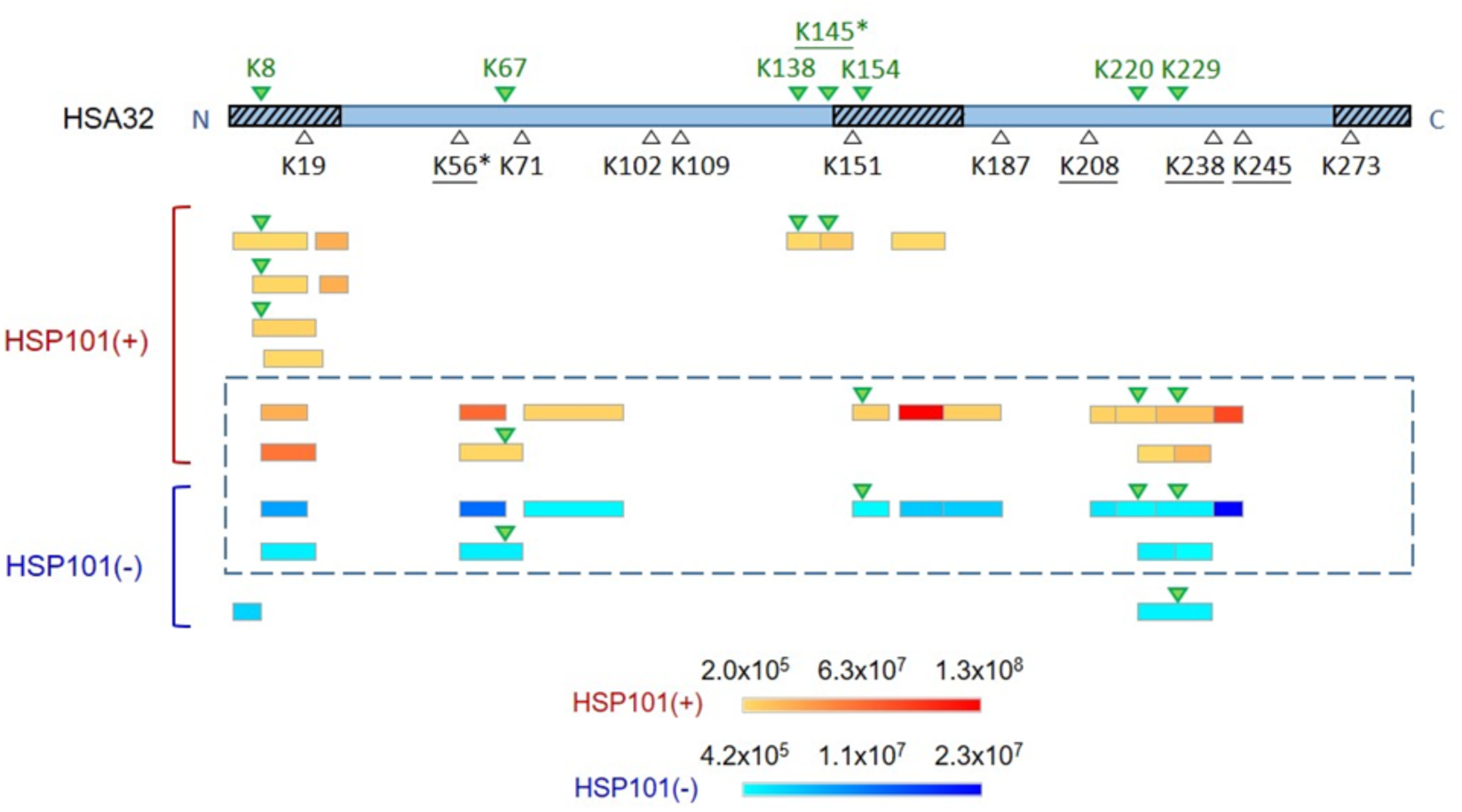
Differential biotinylation of HSA32 by TurboID in the presence and absence of HSP101. The schematic representation of the Arabidopsis HSA32 protein is shown at the top. The three predicted intrinsically disordered regions are marked by hatched boxes at the N-terminus, middle, and C-terminus. Green triangles denote the biotinylated lysines, while white ones are unmodified or non-detected. Stars and underlines indicate the embedded lysine residues in the predicted 3-D structures of HSA32-HA-TurboID and HSA32 trimer, respectively. The identified unique peptides are displayed by colored boxes according to their abundance, from yellow to red for samples with HSP101 (+) and from azure to blue for those without HSP101 (-). Peptides within the dashed box are those commonly identified in the presence or absence of HSP101. N: N-terminus; C: C-terminus; K: lysine.

We also identified the biotinylation sites in the potential interactors of HSA32. Five lysine residues (K871, K872, K884, K893, and K894) at the C-terminus of HSP101 were biotinylated (Supplemental Figure S6C). A single biotinylated lysine residue (K258) was detected on CCT2 (Supplemental Figure S6D).

### HSA32 is an aggregation-prone protein

The protein structure prediction program AlphaFold2 predicted Arabidopsis HSA32 as a TIM-barrel protein, which contains the typical eight alternating β-strands and α-helices with very high pLDDT confidence scores (Figure 7A). However, low pLDDT scores were assigned to three regions at the N- and C-terminus and between the fifth β-strand and fifth α-helix, which coincide with the intrinsic disorder regions (IDRs) predicted by DISOPRED3 (Figure 7B). The predicted 3D structure of HSA32 resembles its homolog in archaea, ComA, determined by x-ray cystography (Wise et al., 2003). A trimeric conformation of HSA32 was predicted by GalaxyHomomer that employs *ab initio* docking of the monomeric structure of HSA32 (Baek et al., 2017), the same as the ComA protein.

**Figure 7.**
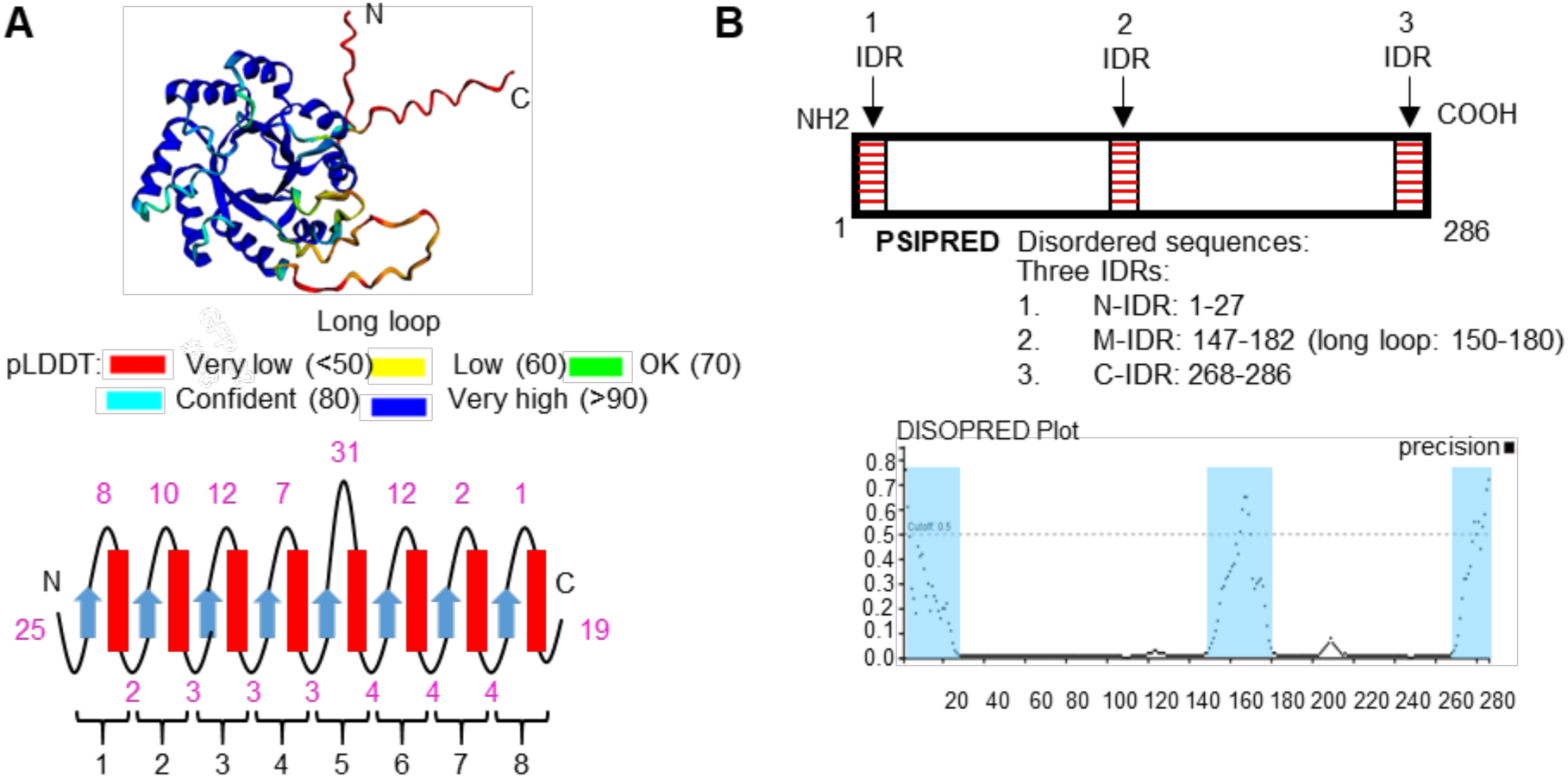
HSA32 is predicted as a TIM barrel protein with IDRs. **A)** The machine-learning algorithm AlphaFold2 was employed to predict the structure of HSA32. The TIM barrel protein HSA32 exhibited unstructured regions at the N- and C-terminus and the long loop linking the fifth β-strand and fifth α-helix. **B)** The three intrinsically disordered regions (IDRs) were predicted on HSA32 using the IDR prediction workbench PSIPRED DISOPRED3. These IDRs were located on the N-terminus, middle domain, and C-terminus of HSA32.

Since the stability of HSA32 depends on HSP101, a protein disaggregase, HSA32 is likely prone to aggregate. Although the HSP101 homolog in yeast is known for prion propagation (Chernoff et al., 1995), we confirmed that HSA32 does not contain a putative Q/N rich prion forming domain with the assistance of a computational method (Zambrano et al., 2015). However, the prediction of the aggregation propensity of HSA32 using AGGRESCAN3D (A3D) based on the AlphaFold2-predicted structure displayed elevated A3D scores at the regions that coincided with the IDRs (Supplemental Figure S7A).

For mass protein production and *in vitro* characterization, the recombinant HSA32 fused to a histidine tag, HSA32-His_6_, was overexpressed in *E. coli*, but it was predominantly found in the fraction of inclusion bodies (Supplemental Figure S7B). Co-expression with the recombinant HSP101 only slightly increased the solubility of HSA32-His_6_ (Supplemental Figure S7B), suggesting that HSP101 is insufficient in forging the solubilization of HSA32 in *E. coli*. The insoluble nature of the recombinant HSA32 in the *E. coli* cells is very different from the case in Arabidopsis seedlings, in which we showed that HSA32 is a soluble cytosolic protein (Figure 3A). We confirmed that HSA32 was detected in the soluble fractions of the crude extracts of both *hsp101* and wild type plants (Supplemental Figure S7C), suggesting that HSP101 does not noticeably affect the solubility of HSA32 *in planta*.

In order to characterize HSA32 extracted from plant cells, we aimed to purify the HSA32 with GFP-tag from the crude extract of the Arabidopsis transgenic plants, *hsa32* (*HSA32-GFP*) and *hsp101 hsa32* (*HSA32-GFP*). The HSA32-GFP proteins could be maximally precipitated at 30-40% ammonium sulfate saturation levels from the crude extracts of *hsa32* (*HSA32-GFP*) and *hsp101 hsa32* (*HSA32-GFP*) seedlings (Supplemental Figure S7D). Intriguingly, the precipitated HSA32-GFP could not be resolubilized by the extraction buffer (data not shown).

## DISCUSSION

Heat acclimation serves as a model system in studying plant stress memories, and HSA32 was the first gene implicated explicitly in HAM (Charng et al., 2006; Bäurle, 2016). Expression of *HSA32* is positively and negatively regulated at the transcriptional level by transcription factors (Charng et al., 2023). The expression of the HSA32 protein is also positively regulated by HSP101 (Wu et al., 2013; Lin et al., 2014), but the underpinning mechanism is unclear. This study showed that HSA32 is regulated post-translationally by HSP101 through modulating its stability. Hence, a positive feedback loop between HSP101 and HSA32 is formed by preventing each other from degradation after HS induces the proteins. We see this feedback loop as one of the core operations of HAM. With HSFA1s, this feedback loop is considered to form a regulatory circuit, which is a part of the complex regulatory networks of heat acclimation response (Charng et al., 2023). Thus, to understand how plants remember their heat stress experience, it is essential to unveil the mechanisms underpinning the core operations of the plant stress memory. Here, we provide a better insight into the interplay between HSA32 and HSP101 by performing multilineage experiments.

Our study has shown that the HSP101-HSA32 feedback loop, which occurs during the memory phase following acclimation treatments at mild (32°C) and extreme (42°C) temperatures, can serve as a core operation for heat stress memory across a wide range of elevated temperatures. However, the differential expression patterns induced by varied high temperatures at the beginning of the memory phase may affect the initial ratio of HSP101/HSA32, potentially influencing the duration of AT. The causes of these different patterns remain to be elucidated in future efforts, suggesting exciting avenues for further research in this field.

Besides induction by heat, HSP101 and HSA32 also accumulate in mature and germinating seeds in Arabidopsis (Supplemental Figure S1D) and rice (Lin et al., 2014). Our results indicate that the interplay between HSP101 and HSA32 can occur without heat treatment in the germinating seeds and very young seedlings (Supplemental Figure S1D), indicating that the factors required for the interplay already exist without being induced or activated by heat at these stages. Despite the HSP101-dependent accumulation of HSA32 in the mature seeds, HSP101 is stable without HSA32. The different effects of HSA32 on HSP101 stability in seeds and seedlings may be due to the different proteostasis controls associated with the developmental stages (Yu and Hua, 2022), which awaits to be examined.

In the interplay relation, how HSP101 positively regulates the accumulation of HSA32 is the focal point of this study. Previously, it was proposed that HSP101 positively regulates the translation of HSA32 (Wu et al., 2013). However, HSA32 could be induced at 32°C in *hsp101* to nearly wild type level at 15 to 30 min into the memory phase (Figure 1B), indicating that HSP101 is dispensable for the translation of HSA32. This observation agrees with a transcriptomic analysis that the *HSA32* mRNA levels associated with polysomes were about equal in wild type and *hsp101* during the memory phase (Merret et al., 2017).

By examining the *in vivo* decay rate of HSA32 using the translation inhibitor CHX applied at the early memory phase, we noticed that HSA32 degrades faster in *hsp101* than in wild type (Figure 1, B and C). Our data further reveal that HSP101 stabilizes the HSA32 protein by preventing its degradation mediated by the proteasome pathway based on the pharmacological results of the proteasome inhibitor BTZ (Figure 2D). Intriguingly, MG132, the most frequently used proteasome inhibitor, failed to prevent the degradation of HSA32 in *hsp101* (Supplemental Figure S2A). The differential effects on HSA32 degradation between MG132 (a peptide aldehyde) and BTZ (a dipeptide boronic acid) may be due to their varying selectivity and specificity in inhibiting the proteasome. Noteworthy, the 20S proteasome exhibits three catalytic activities: chymotrypsin-like (CT-L), trypsin-like (T-L), and peptidyl glutamyl peptide hydrolyzing (PGPH) are mediated by three distinct catalytic β5, β2, and β1 subunits, respectively (Groll et al., 1999). In mammalian studies, it was shown that BTZ completely inhibited CT-L (β5) and PGPH (β1) activities, whereas MG132 displayed weak inhibition of these catalytic activities (Crawford et al., 2006; Kloß et al., 2010). Furthermore, it has been reported that the efficacy of the inhibitors varies with the nature of the protein substrate (Kisselev et al., 2006). Hence, BTZ likely functions as a more specific and potent inhibitor for HSA32 degradation than MG132. Similar situations were observed for the degradation of HSP101. MG132 cannot block the degradation of HSP101 (Wu et al., 2013; McLoughlin et al., 2019), but BTZ can (Figure 2E).

Macroautophagy was shown to reset the stress memory by slowly degrading heat-induced HSPs during the memory phase (Sedaghatmehr et al., 2019), including HSP101 and sHSP-CI confirmed in this study (Figure 2A). Thus, proteasome and autophagy share an overlapped duty in shaping HAM. We further showed that autophagy participates in HSA32 degradation in the presence of HSP101 but not in its absence (Figure 2, A and B), suggesting that autophagy is responsible for the slow clearance of HSA32 that HSP101 has stabilized. Without HSP101, a significant fraction of HSA32 is rapidly degraded by proteasome, but not by autophagy, which is consistent with earlier observations that the autophagy inhibitor, E-64d, did not block the degradation of HSA32 in *hsp101* (Wu et al., 2013). In this work, we show that proteasomes also contribute to the degradation of sHSP-CI during the memory phase, which manifests to degrade faster than HSP101 and HSA32 and can be substantially rescued by inhibiting proteasome activity with BTZ (Figure 2D). Interestingly, the role of the proteasome in degrading HSA32 and HSP101 becomes more pronounced in *hsp101* and *hsa32*, respectively, than in wild type (Figure 2, D and E), suggesting that the interplay between HSP101 and HSA32 provides mutual protection to each other against proteasome degradation. This protection should be specific to a certain extent, as the proteasome-mediated degradation of sHSP-CI was not affected by HSP101 (Figure 2D).

The rapid degradation of HSA32 by the proteasome in *hsp101* raised the question of whether this process involves polyubiquitination, a prerequisite for the degradation of many proteins via the 26 proteasome pathway (Smalle and Vierstra, 2004). Our answer is no based on the results of different experiments. First, we noticed an increased accumulation of polyubiquitinated proteins in *hsp101*, consistent with the recent report (McLoughlin et al., 2019). However, polyubiquitinated HSA32 could not be detected (Supplemental Figure S2F). Secondly, inhibiting E1 activity or disrupting the 19S regulatory particle of the proteasome did not prevent HSA32 degradation (Supplemental Figure S2, G and I). Thirdly, E3 ligases could not be identified by HSA32-HA-TurboID with or without HSP101 (Figure 4, C and D), despite the successful cases of the identification of E3 ligases by similar approach (Zhang et al., 2019; Lee et al., 2023). Examples have been reported that proteins can be degraded by proteasome without being polyubiquitinated (Ben-Nissan and Sharon, 2014). Whether it is also the case for HSA32 remains to be investigated.

How does HSP101 confer protection to HSA32? There are several clues to the answer to the question. First, HSP101’s disaggregase function may play an essential role in the protection. It was shown that the point mutations that compromise HPS101 function led to a faster degradation of HSA32 during the memory phase (Wu et al., 2013). Several lines of evidence indicate that HSA32 is prone to aggregate, making it an ideal target for HSP101. HSA32 was shown to interact with itself, even in the presence and absence of HSP101 (Supplemental Table S, 2 and 4). It is worth mentioning that TIM-barrel proteins have the ability to self-associate to form reversible aggregates (Rodríguez-Bolaños et al., 2020). We hypothesize that after folding, TIM-barrel HSA32 proteins might be self-associated to form aggregates. Although the predicted structural basis for the aggregation propensity of HSA32 remains to be validated (Supplemental Figure S7A), our results point out its insolubility following ammonium precipitation and when heterologously expressed in *E. coli*. HSP101 likely protects HSA32 by preventing it from aggregation, which could be toxic to the cells and susceptible to fast degradation. The action of HSP101 on the HSA32 aggregates may resemble that of yeast HSP104 on preamyloid oligomers, phase-transitioned gels, disordered aggregates, amyloids, or prions ((Shorter and Southworth, 2019). However, the exact action mode of the disaggregase depends on the nature of HSA32 aggregates, which remains to be determined.

The action of a protein disaggregase on its substrate requires a direct interaction based on the mechanism of HSP104 in yeast and ClpB in *E. coli* (Doyle and Wickner, 2009). The results of the proximity labeling experiments support that HSP101 directly interacts with HSA32 because HSP101 was the most prominent target of HSA32-HA-TurboID (Figure 4C and Supplement Table S2). Unfortunately, other traditional methods, such as Co-IP, Y2H, and BiFC, could not validate this finding, most likely due to either the transient interaction between the two proteins or the tendency of HSA32 to aggregate. Several studies showed that proximity labeling-based methods outperform Co-IP or Y2H for detecting transient interactions (Roux et al., 2012; Arora et al., 2020; Kim et al., 2023). Moreover, attaching a tag to HSP101 might weaken or disrupt the interaction. Indeed, we observed that a tag at the N- or C-terminus of HSP101 hampered the interaction between HSP101 and HSA32 (Figure 5, D and E).

In addition to HSP101, three subunits of the cytosolic chaperonin TRiC/CCT complex (CCT2, CCT3, and CCT8), AtJ3 (a co-chaperone of HSP70), and P23-1 (a co-chaperone of HSP90) were identified as potential interacting proteins of HSA32. Their interaction with HSA32 is independent of HSP101 (Figure 4, C and D). In mammalian cells, TRiC/CCT has been shown to efficiently fold newly synthesized proteins ranging from ∼30 to 60 kDa (Frydman et al., 1994; Thulasiraman et al., 1999). HSP70 and HSP90 facilitate the delivery of the partially folded substrates to the TRiC/CCT complex for appropriate folding (Melville et al., 2003; Cuéllar et al., 2008). Moreover, the clients of TRiC/CCT were predicted to be slow-folding and aggregation-prone (Yam et al., 2008). Hence, we hypothesize that TRiC/CCT, with the assistance of HSP70-AtJ3 and HSP90-P23-1, plays a role in the folding of newly synthesized aggregation-prone HSA32, similar to the mechanism observed in the mammalian system. HSP101 might not be needed for this folding process as HSA32 could interact with TRiC/CCT without HSP101. Thus, the folded HSA32 tends to aggregate, which HSP101 reverses. This notion is supported by the differential biotinylation sites on HSA32, *i.e.*, the pattern of biotinylation on HSA32 differed depending on the presence or absence of HSP101 (Figure 6 and Supplemental Table S6). These findings lead us to propose two possible explanations: 1) HSP101 modifies the conformation of HSA32 so that these lysine residues become accessible to TurboID; 2) these lysine residues were blocked by other interacting protein (s) in the absence of HSP101. The latter possibility aligns with our findings of distinct interacting partners of HSA32 depending on the presence or absence of HSP101 (Figure 4, C and D). To our knowledge, this is the first study demonstrating that prey protein (HSP101) can influence the biotinylation pattern of a bait protein (HSA32) fused to TurboID.

The localization studies revealed that the newly synthesized HSA32-GFP formed foci in response to mild heat stress (32°C), and these foci exhibited liquid-like properties and gradually dispersed over time (Figure 3C). A question arises as to whether these HSA32 foci could be stress granules (SGs). Two lines of evidence suggest that they are not canonical SGs. First, SGs are formed only above 35°C (Hamada et al., 2018). We showed that the SG marker PAB2-RFP condensated at 37°C but failed to do so at 32°C (Supplemental Figure S3H). Next, several studies have shown that SGs exhibit fusion behavior of the droplets in mammalian, yeast, and plant cells (Kroschwald et al., 2015; Tong et al., 2022; Zhu et al., 2022). However, the HSA32 foci gradually disappeared over time and did not undergo a fusion event during the recovery phase (Supplemental Figure S3G). These observations suggest that HSA32 foci might represent a type of condensate different from the SGs. In addition, it has been demonstrated that many IDR-containing proteins that degraded by 20S proteasome were tightly linked with phase-separated condensates (Myers et al., 2018). We observed that HSA32 foci formation emerged after 30 min of heat stress (Supplemental Figure S3D), suggesting that the foci are associated with the folding process following HS. The pre-existing HSA32-GFP does not form foci in response to HS (Figure 3G, the sample with CHX treatment). Hence, we hypothesized that the nascent HSA32 is prone to aggregate and form foci through LLPS following heat stress. In Arabidopsis, the protein DXPS formed punctate spots following heat treatment, representing aggregated DXPS (Yu et al., 2021). In this connection, chaperone-mediated disaggregation was also observed by chaperone sHSP21, preventing aggregation of DXPS for providing thermoprotection in Arabidopsis (Yu et al., 2021). HSP101 (McLoughlin et al., 2019) and AtJ3 (Wang et al., 2021) have been observed to form cytosolic foci following heat acclimation in plants. In yeast, the formation of protein aggregates (foci) is not associated with the chaperone HSP104, but the disassembly of the aggregates depends on the disaggregase function of HSP104 (Cabrera et al., 2020). Our findings align with these observations as HSA32 foci can form without HSP101, but the number of foci significantly decreases when HSP101 is present during recovery periods (Figure 3, C and D).

Finally, we propose a working model to depict the interplay between HSA32 and HSP101 (Figure 8). After HS induction, HSP70/HSP90 and TRiC/CCT complexes facilitate the folding of the newly synthesized HSA32 proteins. In the absence of HSP101, folded HSA32 proteins tend to form aggregates, which are vulnerable to degradation by proteasome in a polyubiquitin-independent manner. HSP101 can recognize, bind, and cleave the HSA32 aggregates, rendering them disaggregated and stable. Subsequently, the stabilized HSA32 prevents the degradation of HSP101 via a yet unknown enzymatic reaction. As a result, acquired thermotolerance is prolonged. The involvement of protein aggregation in plant stress memory is striking, as protein aggregation has been a feature in the memories of animals and yeast (Santos and Ventura, 2021). There might be a convergent function of protein aggregation in memory during evolution. Our findings here provide an example of plants. Based on the observed role of HSP101 and HSA32 in HAM, it is suggested that these genes could be promising targets to develop HS tolerant crop plants, opening new avenues for research in agricultural biotechnology.

**Figure 8.**
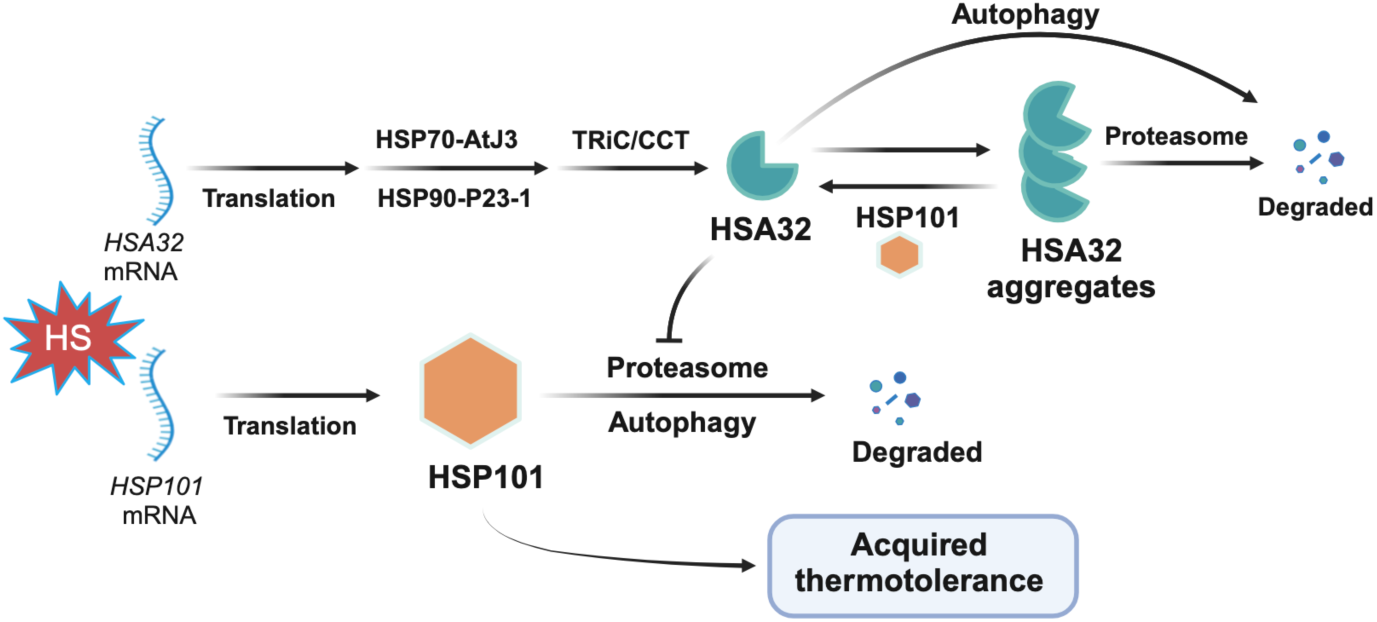
The proposed working model of the HSP101-HSA32 module. The proposed model shows heat acclimation upregulates HSA32 and HSP101 mRNAs and protein synthesis. Newly synthesized HSA32 proteins are folded with the assistance of HSP70-AtJ3, HSP90-p23-1, and TRiC/CCT chaperonin. After that, the folded HSA32 remains aggregation-prone and readily forms aggregates. HSP101 directly interacts with the HSA32 aggregates and triggers their disaggregation. Without HSP101, the HSA32 aggregates are susceptible to proteasome-mediated faster degradation. In the presence of HSP101, HSA32 can also be degraded slowly via the autophagy pathway. On the other hand, HSA32 prevents the degradation of HSP101. As a result, acquired thermotolerance is prolonged. Without HSA32, HSP101 is rapidly degraded via proteasome and autophagy pathways, leading to faster decay of acquired thermotolerance.

## MATERIALS AND METHODS

### Plants materials and growth conditions

*Arabidopsis thaliana* T-DNA KO mutants, *hsp101* (SALK_066374), *hsa32* (GABI-Kat-268A08), and double mutant *hsp101 hsa32* were described previously (Wu et al., 2013). The double mutant *hsp101 atg5* was generated through crossing between *atg5* (SALK_020601) and *hsp101*. The transgenic line expressing PAB2-RFP was reported previously (Merret et al., 2017). All T-DNA insertion mutations were confirmed by PCR analysis with gene-specific primers, and homozygous lines were selected. For seedlings, Arabidopsis seeds were sterilized and sown on Murashige and Skoog (MS) medium plates containing (24 mL of 0.5x MS with 1% sucrose and 0.8% agar) and stratified for 3 d at 4°C and then transferred to growth chamber at 22°C and 16 h of light (120 µmol m^-2^ s^-1^).

### Thermotolerance assays

A water bath was utilized to conduct the LAT and SAT assays for heat acclimation treatments (Yeh et al., 2012). For the LAT assay, 3-d-old seedlings were exposed to the first HS at 32°C for 1 h and recovered for 1 d, and then a second HS was performed at 44°C for 45 min and recovered for 10 d when images were taken. For the SAT assay, 4-d-old seedlings were subjected to the first HS at 32°C for 1 h and recovered for 2 h, and then a second HS was carried out at 44°C for 50 min and recovered for 10 d when images were captured.

### Immunoblotting

The proteins were separated by SDS-PAGE (NuPAGE 4–12% Bis-Tris gel, Invitrogen) and transferred onto a nitrocellulose membrane for antibody probing. Blots were probed with primary antibodies for overnight incubation at 4°C, followed by 1 h incubation at room temperature with secondary antibodies. ImageJ software (National Institutes of Health) was applied to quantify the intensity of the protein bands. The antibodies against HSA32 and HSP101 were described previously (Chi et al., 2009). Several antibodies were used for immunoblotting, including rabbit anti-sHSP-CI (Abcam), mouse anti-Tubulin (Sigma), mouse anti-HA (Abcam), rabbit anti-Ub (Millipore Sigma), rabbit anti-GFP (Abcam), SA-HRP (Abcam), mouse anti-H3 (Abcam), rabbit anti-COXII (Agrisera), and mouse anti-His (Novagen).

### Chemical treatments for protein degradation assay

Freshly prepared cycloheximide (50 µM), MG132 (100 µM), bortezomib (50 µM), and PYR-41 (50 µM) were applied to Arabidopsis seedlings (350 µL solution per 50 seedlings) on the plate at 15 min recovery period after HS treatment and collected samples at the indicated time. The chemicals were dissolved in 100% DMSO, and the working solutions were prepared in 0.1% DMSO solution, which served as negative control.

### TUBEs pull-down assay

The TUBE assay was carried out following previous procedures with modifications (Lee et al., 2020). Leaf tissues were finely ground, and total plant proteins were extracted using cell lysis buffer (50 mM Tris-HCl, pH 7.5, 150 mM NaCl, 1mM EDTA, 1% NP-40, 10% glycerol, 1 mM PMSF, and 1% of plant protease inhibitor cocktail, and 50 µM of PR-619 deubiquitinating inhibitor). The protein solution was incubated at 4°C for 30 min with rotation and then subjected to centrifuge at 14,000g for 15 min at 4°C. The supernatant was collected, and 3.5 mg of proteins were precleared with agarose beads for 1 h at 4°C through rotation. The precleared samples were then distributed into two fresh tubes and incubated with either the TUBE-conjugated agarose (LifeSensors) or control agarose beads at 4°C with rotation for 16 h. After incubation, the beads were washed five times with TBS-T buffer (20 mM Tris, pH 8.0, 150 mM NaCl, 0.1% Tween-20), and total bound proteins were eluted by 1.5× Laemmli buffer after incubating at 65°C for 10 min. The pull-down ubiquitinated proteins were detected by immunoblotting using an anti-ubiquitin antibody (Millipore Sigma, 05-1307, clone Apu2) at a concentration of 1:1000. The ubiquitination level of HA-HSA32 was examined employing an anti-HA antibody at a concentration of 1:1000 (Sigma-Aldrich, H3663, clone HA-7). The experiment was repeated three times with similar outcomes.

### Subcellular fractionation analysis

Subcellular compartments of heat-acclimated seedlings were isolated from wild type and *hsa32* (*GFP-HSA32)* seedlings using differential ultra-centrifugation of their cell lysates. The 5-d-old seedlings were subjected to heat treatment at 37°C for 1 h and collected samples at 16 h recovery period. Four distinct fractions were collected and analyzed: nuclear fraction (NF), mitochondrial fraction (MF), microsomal fraction (MS), and cytosolic soluble fraction (CS). For nuclear faction, centrifuge was performed at 10 min at 600 x g and collected pellet as crude nuclear faction and supernatant used for further centrifuge at 30 min at 10K x g and collected pellet as mitochondrial-rich fraction pellet. Next, the supernatant was used for further centrifuge at 2 h at 100K x g and collected pellet as total microsomal fraction and supernatant represented cytosolic soluble proteins. Anti-histone3 (H3), COXII, and proteasome antibodies were used as a marker for nuclear, mitochondrial, and cytosolic soluble fractions, respectively.

### Confocal microscopy analysis

Vertically grown 4-day-old seedlings were used for confocal microscope (Zeiss LSM 780) to observe protein subcellular localization. For heat treatment at 32°C for 1 h, the plates with seedlings were placed vertically into a water bath and immediately observed under a confocal microscope. The number of foci was quantified with Image J software. In chemical treatments, cycloheximide (50 µM) and bortezomib (50 µM) solution were added to the plate with seedlings before 1 h of heat treatment and kept in a growth chamber for incubation. Subsequently, the plates were transferred in the water bath at 32°C for 1 h, and then the samples were immediately observed under a confocal microscope. Fluorescence recovery after photobleaching (FRAP) assay was executed as previously described (Tong et al., 2022). A laser intensity of 100% with a wavelength of 488nm was applied on one speckle or focus to induce bleaching, and the recovery process was monitored using 2% of the maximum excitation laser power. Images were captured before bleaching, bleaching, and time-lapse post-bleaching for the indicated periods using a 63 x oil immersion objective. For the stress granule test, a transgenic plant with stress granule marker gene Wt (*PAB2-RFP*) was crossed with *hsa32* (*HSA32-GFP*) to stack the transgenes.

### Generation of transgenic lines expressing HA- and GFP-tagged HSA32

The expression of all the HSA32 complementation lines in the Arabidopsis *hsa32-1* background was controlled by a constitutive 35S promoter. For the expression of HA-HSA32, 35S promoter, 3X HA, HSA32 CDS, and NOS terminator were amplified by PCR and cloned into pCR8/GW/TOPO TA vector (Invitrogen). The constructs were sequenced to confirm no missense or nonsense mutation in the coding region. The vectors were then subcloned into pCAMBIA1390 by restriction enzymes and T4 DNA ligase (Promega) according to the manufacturer’s protocol. HSA32 and GFP were cloned into pCR8/GW/TOPO TA vector and then subcloned into pMDC45 and pB2GW7-0, respectively, by LR reaction for the expression of GFP-HSA32 and HSA32-GFP. The transgenic lines were generated by the floral dip method.

### Generation of HA fused HSP101 transgenic lines

3XHA-HSP101 fusion proteins were expressed under the control of the *A. thaliana HSP101* (*AtHSP101*) promoter in *hsp101*. The recombinant DNA fragments of *AtHSP101p::3X HA*-*HSP101* were amplified by overlap extension PCR. The PCR product was purified, cloned into pCR8/GW/TOPO TA vector (Invitrogen), and sequenced. The vectors were then subcloned into pCAMBIA1390 by restriction enzymes and T4 DNA ligase (Promega) according to the manufacturer’s protocol.

### TurboID constructs and generation of transgenic lines

The NEBuilder HiFi DNA assembly master mix (New England Biolabs) was used for generating three constructs, where the 3HA-TurboID (Addgene plasmid# 107171) was fused to the N- or C-terminus of HSA32. Additionally, a construct with 3HA-TurboID was generated to serve as a control. All three constructs were developed under the control of the *HSA32* promoter (1 kb upstream of the start codon). The ligated products were cloned into the pCAMBIA1390 vector, and transformation was performed on competent *E. coli* (TOP10). All transgenes were confirmed through sequencing. Agrobacterium strain GV3101 harboring different constructs was transformed into the *hsa32* mutant by floral dip. Homozygous T3 seeds were used for all experiments. Selected T3 homozygous transgenic lines were crossed with a double mutant *hsp101 hsa32* to generate transgenic lines with double mutant background.

### Biotinylation of proteins by TurboID fusion proteins *in vivo*

The preparation of biotin solution, protein extraction, and streptavidin beads preparation were performed as described previously (Khan et al., 2018). Following heat treatment, 4-d-old transgenic seedlings were submerged in beakers containing biotin solution and incubated at room temperature for the indicated periods. After protein extraction, SDS-PAGE gels were run and probed using Streptavidin-horseradish peroxidase conjugate SA-HRP (Abcam, ab7403) at a concentration of 1:10,000 and HA antibody to examine the biotinylated proteins.

### Agroinfiltration

The *Agrobacterium tumefaciens* GV3101 cells carrying various constructs were infiltrated into *N. benthamiana* leaves, as described previously (Zhang et al., 2019). The cultured agrobacteria were initially inoculated into LB broth with 50 μg/mL kanamycin, 10 mM MES, and 20 µM acetosyringone in a 14-mL Falcon culture tube with shaking at 28°C overnight. Subsequently, the agrobacteria were centrifuged at 3,000 x g for 10 min. The pellet was resuspended to OD_600_ = 1 in the suspension solution containing 10 mM MgCl_2_, 10 mM MES, and 100 µM acetosyringone. For co-infiltration, different agrobacteria cultures were mixed in equal proportion (1:1) and then infiltrated into the leaves of 3- to 4-wk-old *N. benthamiana* plants. Three days post agroinfiltration, a solution of 200 μM biotin was infiltrated into the same sectors of the leaves. The infiltrated leaves were cut at the base of the petiole with the leaf vein removed and frozen in liquid nitrogen for protein extraction.

### Enrichment of biotinylated proteins for LC-MS/MS analysis

Following protein extraction, the solution was run through the PD-10 desalting columns (Sephadex G-25 Media) according to the manufacturer’s instructions to remove excess biotin. Then, protein concentration was determined. Around 2.5 mg of proteins were incubated with 50 µL of the washed beads at 4°C on a rotator for 16 h. The beads were subsequently centrifuged for 2 min at 500g at 4°C, and a magnetic rack was used to isolate the beads. The beads were suspended with 1 mL of extraction buffer without detergent, followed by centrifugation at 500g for 30 sec at 4 °C to collect the beads. The beads were washed with SDS buffer (2% SDS, 20 mM Tris-HCl, pH 7.5), protein extraction buffer without detergent, and 50 mM ammonium bicarbonate (ABC, pH 8.0), and finally, the beads were resuspended in 1 mL of 50 mM ABC for LC-MS/MS analysis.

Samples were lysed in 5% (v/v) SDS in 50 mM triethylammonium bicarbonate (TEAB). The solution was transferred into a 1.7-mL tube and sonicated 10 times for 10 s each. The tube was centrifuged at 16,000 g at 4°C for 20 min, and the supernatant was collected. The protein amount was determined by bicinchoninic acid assay (Thermo Fisher Scientific). Protein digestion in the S-Trap microcolumn was performed according to the previous protocol with some modifications (Chen et al., 2023). Briefly, proteins in the lysis buffer were reduced and alkylated using 10 mM Tris (2-carboxyethyl) phosphine hydrochloride (TCEP) and 40 mM 2-chloroacetamide (CAA) at 45°C for 15 min. A final concentration of 5.5% (v/v) phosphoric acid (PA) followed by a six-fold volume of binding buffer (90%, v/v, methanol in 100 mM TEAB) was next added to the protein solution. After gentle vortexing, the solution was loaded into an S-Trap microcolumn. The solution was removed by spinning the column at 4,000 g for 1 min. The column was washed with 150 μL binding buffer three times. Finally, 20 μL of digestion solution (1 unit Lys-C and 1 μg trypsin in 50 mM TEAB) was added to the column and incubated at 47°C for 2 h. Each digested peptide was eluted from the S-Trap micro column using 40 μL of three buffers consecutively: (1) 50 mM TEAB, (2) 0.2% (v/v) FA in H_2_O, and (3) 50% (v/v) ACN. Elution solutions were collected in a tube and dried in a vacuum.

Peptides were suspended in 40 µL 0.1% (v/v) formic acid (FA) with 3% (v/v) ACN, and 4 µL of the sample was injected into an UltiMate3000 UHPLC system coupled with an Orbitrap Q-Exactive (Thermo Fisher Scientific). Buffer A was 0.1% (v/v) FA, and buffer B was 0.1% (v/v) FA in 100% ACN. The LC gradient of 60 min was established as follows: 5 min from 3% to 5% buffer B, 55 min from 5% to 25% buffer B, 2 min from 25% to 85% buffer B, 5 min buffer B, 1 min from 85% to 3% buffer, and 22 min 3% buffer B. Peptides were separated on a column with a Waters nanoEase M/Z Peptide CSH C18 25mm column heater set at 50°C. The mass spectrometer was operated in data-dependent acquisition mode, in which a full MS scan (m/z 350-1600; resolution: 70,000) was performed. Data were acquired in the Orbitrap with a resolution of 17,500 [normalized collision energy (NCE): 27%; max injection time: 100ms; isolation window: 2.0 m/z; dynamic exclusion: 20s].

The MS raw files were searched against the TAIR10 database (35,386 entries, release date: 2019-07-11) using Sequest and Mascot search engines in Proteome Discoverer 2.5. Precursor mass tolerance was set to ±10 ppm, and fragment mass tolerance was set to 0.02 Da. Protein digestion was set to trypsin (after KR/-) up to two missed cleavages. The fixed modification was set as carbamidomethyl (C), and variable modifications were set as oxidation (M) and acetylation (protein N-term). The FDR for peptide-spectrum match (PSM), ion, peptide, and protein level was set at 1%. The protein abundances from three biological replicates were uploaded to Perseus software 1.6.5.0 (Tyanova et al., 2016). The abundance values were transformed to log2 format, normalized by subtraction of the median, and filtered for the valid values in each group. The missing values were replaced from a normal distribution using a standard setting in Perseus, with a width of 0.3 and a down-shift of 1.8. To determine the significantly enriched proteins in HSA32-HA-TurboID samples compared to those in HA-TurboID ones, Student’s T-test was applied and visualized in scatter plots with a cutoff of 1.0 set for the log2(fold-enrichment) and 1.3 for the –log(*p*-value).

### Expression of recombinant proteins in *E. coli*

To express HSA32-His6and HSP101 in *E. coli*, two plasmids (pET-24a(+)-AtHSA32-His, pET-32a(+)-AtHSP101) were individually transformed or co-transformed into chemically competent BL21 (DE3) cells by heat-shocking. The transformed cells were selected on an LB agar medium containing kanamycin (50 μM/mL), ampicillin (100 μM/mL), or both. For the induction of the recombinant proteins, 0.1 M IPTG was added to the *E. coli* cultures grown at 22 or 37 °C in LB medium with antibiotics. The *E. coli* cells were suspended in the extraction buffer (50 mM Tris-base (pH 7.5), 500 mM NaCl, 2 mM EDTA, 0.1% Triton X-100, 0.1 mg/mL lysozyme, and 1% protease inhibitor cocktail) and vortexed vigorously. The cell suspension was sonicated using an S-4000 sonicator (MISONIX) at 70 amplitude and 30% duty cycle for 10 min. Cell lysates were centrifuged at 17,000 x g for 15 min at 4°C, and the supernatants were then transferred to fresh tubes (the soluble fraction). The pellets were washed six times with 1 mL of washing buffer (50 mM Tris pH 7.5, 500 mM NaCl, and 2 mM EDTA) and 0.1 g of silicon carbide by pipetting and vertexing. The washed pellets were dissolved in SDS extraction buffer (50 mM Tris pH 7.5, 500 mM NaCl, 2 mM EDTA, 0.1% Triton X-100, and 2% (w/v) SDS) followed by centrifugation at 17,000 x g for 15 minutes to remove debris. The supernatant represents the insoluble fraction of the recombinant proteins. The soluble and insoluble proteins were subjected to immunoblotting analysis as described earlier.

### Computational tools for prediction analysis

We employed computational tools to predict the structure and properties of HSA32. The protein’s 3D structure of HSA32 was predicted using a machine-learning algorithm, AlphaFold2 (https://alphafold.ebi.ac.uk). Intrinsic disordered (ID) regions were predicted using workbench PSIPRED DISOPRED3 (http://bioinf.cs.ucl.ac.uk/psipred/) (Jones and Cozzetto, 2015). HSA32 aggregation properties were analyzed using AlphaFold predicted structure and were downloaded from MODataBase of Aggrescan3D 2.0 server, and the predicted aggregation scores were obtained (https://biocomp.chem.uw.edu.pl/A3D2/) (Kuriata et al., 2019).

## Acknowledgments

The TurboID plasmid (Addgene plasmid# 107171) is kindly provided by Shu-Hsing Wu (IPMB, Academia Sinica). We thank Chun-Kai Huang (IPMB, Academia Sinica) for guiding the TurboID experiment. The transgenic line expressing NPR1-GFP is kindly supplied by Hsin-Hung Yeh (ABRC, Academia Sinica). We thank Academia Sinica Advanced Optics Microscope Core Facility (AS-CFII-111-208) for microscope imaging technical support. In addition, we thank Ansar Ali (IPMB, Academia Sinica) for assisting in the operation of the confocal microscope. We thank the Academia Sinica DNA Sequence Core Facility (AS-CFII-111-211) of the Institute of Biomedical Sciences of Academia Sinica for providing DNA sequence services.

