## Supplemental Figures for "Stabilization of HSA32, an aggregation-prone protein, by the protein disaggregase HSP101 plays a critical role in maintaining acquired thermotolerance in Arabidopsis"

##### Supplemental Figure S1

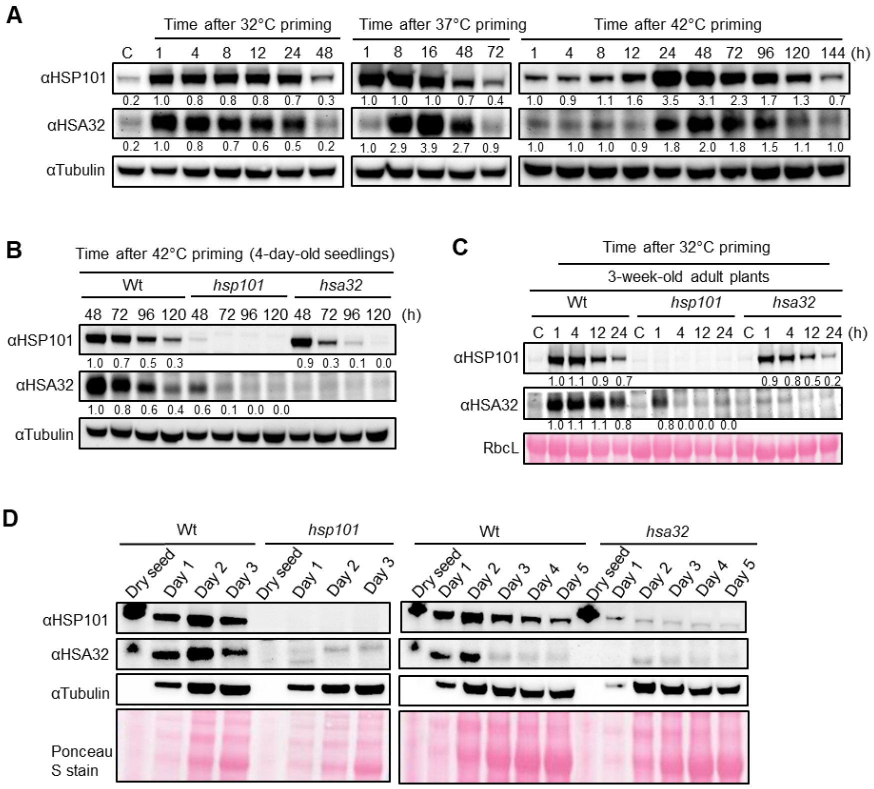


**Supplemental Figure S1. Effect of temperature on induction of HSP101 and HSA32 proteins.** **A)** Wild type (Col-0) seedlings were subjected to three different heat acclimation treatments (32°C, 37°C, and 42°C), recovered for the indicated periods, and examined the protein level of HSP101 and HSA32 using immunoblotting analysis. Total proteins were resolved on SDS-PAGE, and 50 µg protein was loaded in each well. The blots were probed with antibodies against HSP101, HSA32, and Tubulin. ImageJ software was employed to quantify the intensity of the protein bands. **B)** The interplay between HSP101 and HSA32 was studied at 42°C for 30 min in 4-d-old seedlings of wild type, *hsp101*, and *hsa32* mutant plants. The blots were probed with antibodies against HSP101, HSA32, and Tubulin. **C)** The interplay between HSP101 and HSA32 during adult stage. The leaves were collected from 3-wk-old adult plants, kept on the wet tissue in the Petri dish for exposure to heat treatment at 32°C for 1 h, and recovered for indicated time points. The blots were probed with antibodies against HSP101, HSA32, and Tubulin. **D)** Seeds were sterilized and imbibed at 4°C for 3 days in the dark and then transferred to a growth chamber. The samples were collected on the specified days under normal temperature (22°C) to investigate the interplay. Mature, dry seeds were also used after sterilization. The antibodies against HSP101 and HSA32 were used to examine the protein levels, and tubulin and Ponceau stain were used as loading control.

**Supplemental Figure S2**

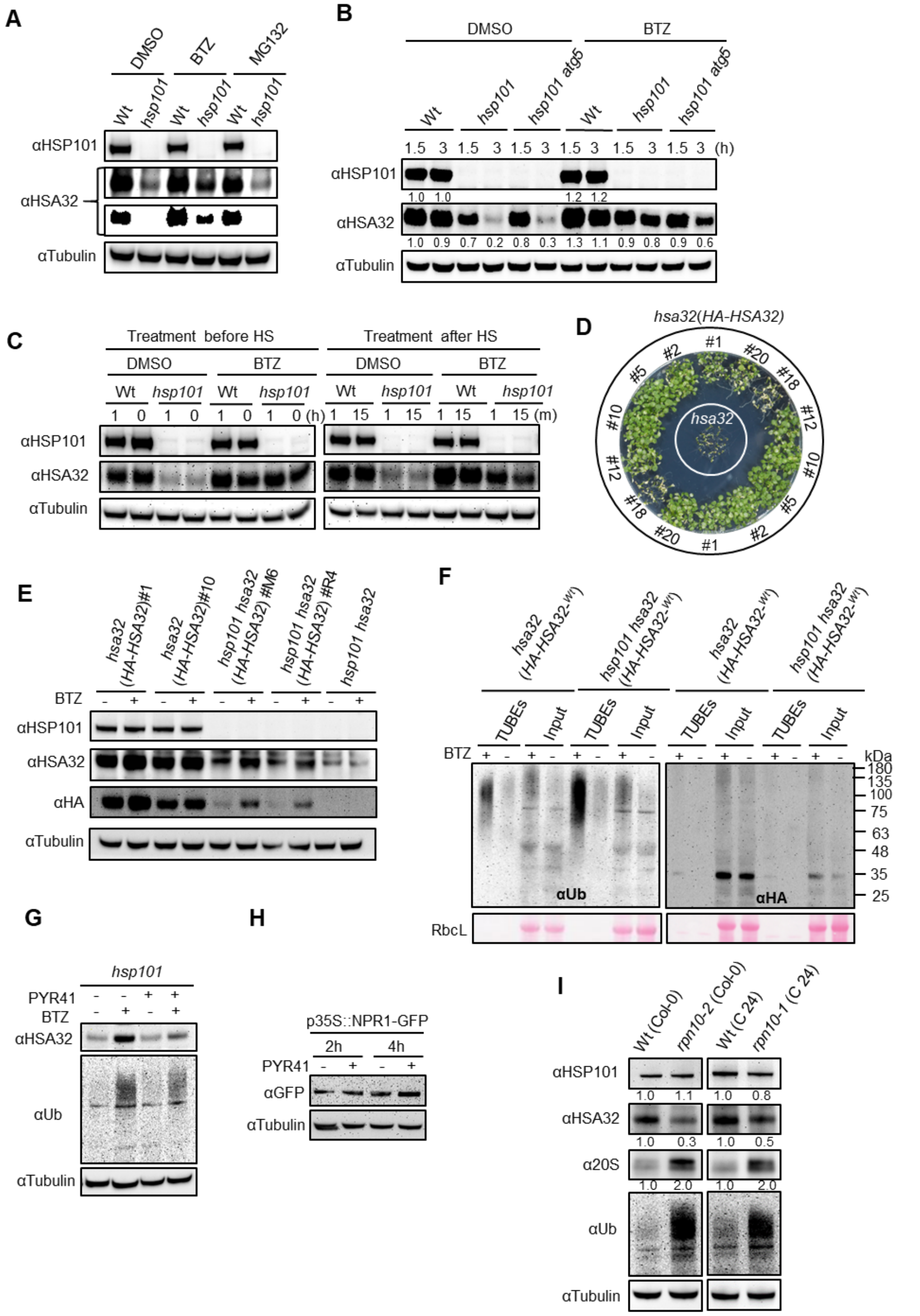


**Supplemental Figure S2. The degradation processes of HSA32. A)** Effect of proteasome inhibitors MG132 and Bortezomib (BTZ) on the degradation of HSA32. DMSO (0.1%), MG132

(100 μM), and BTZ (50 μM) were applied onto 4-d-old seedlings (350 μL solution per 50 seedlings) on the plate at 15 min recovery period following HS at 32°C for 1h and the samples were collected at 2.5 h recovery period. Antibodies against HSP101, HSA32, and Tubulin were used to probe the blots, and tubulin served as loading control. **B)** Effect of BTZ on *hsp101* and *hsp101 atg5*. The seedlings were treated with DMSO or BTZ (50 μM) at 15 min recovery after experiencing HS treatment and recovered indicated time points. The protein levels of HSP101 and HSA32 were examined with an anti-HSA32 antibody. **C)** The comparison of the application timing of BTZ. The DMSO or BTZ was applied one hour (1 h) and immediately before (0 h) heat stress, as well as immediately after heat treatment (1 min) and a 15 min recovery period. All samples were collected at 2.5 h of recovery for immunoblotting. **D)** The LAT phenotypes of the HA-HSA32 transgenic lines in the *hsa32* background were examined for the complementation analysis. **E)** The protein level of HSA32 was assessed in the HA-HSA32 transgenic lines in *hsa32* single and *hsp101 hsa32* double mutant background in the presence or absence of proteasome inhibitor BTZ. BTZ (50 μM) was applied to investigate the inhibitory effect on the degradation of the transgenic lines of HSA32. All transgenic lines were treated with BTZ (+) or DMSO (-) after HS treatment and recovered for 3 h. **F)** Transgenic lines *hsa32* (*HA-HSA32*) and *hsp101 hsa32* (*HA-HSA32*) were treated with bortezomib (+) or DMSO (-) after HS treatment at 32°C for 1 h and kept for recovery 3 h. TUBE resins were used to purify the ubiquitinated proteins, and agarose resins were used to detect nonspecific protein binding that served as input. Anti-Ub and anti-HA antibodies were used to detect total ubiquitinated proteins (left panel) and ubiquitinated HA-tagged HSA32 (right panel), respectively. Molecular mass markers in kilodaltons (kDa) were indicated on the right. **G)** The seedlings of *hsp101* were treated with DMSO (0.1%), PYR41 (50 μM), or bortezomib (50 μM) following HS at 32°C for 1 h, and the samples were collected at 3 h recovery periods. The blots were probed with anti-HSA32, Ub, and tubulin. **H)** The transgenic line p35S: NPR1-GFP was used as a control to evaluate the effectiveness of PYR41. Samples were collected at 2 and 4 h recovery after application of PYR41 and DMSO. **I)** The protein level of HSA32 was assessed in both wild type (Col-0 and C24 ecotypes) and 19S proteasome subunit RNP10 mutants (*rpn10-1* and *rpn10-2*) to examine the involvement of 19S proteasome in HSA32 degradation. Notably, *rpn10-1* mutant is in the C24 ecotype background, while *rpn10-2* is in the Col-0 ecotype background. The blots were probed with antibodies against HSP101, HSA32, 20S proteasome, ubiquitin, and tubulin. Each well contained 50 μg protein, and tubulin was used as a loading control for all immunoblotting assays.

**Supplemental Figure S3**

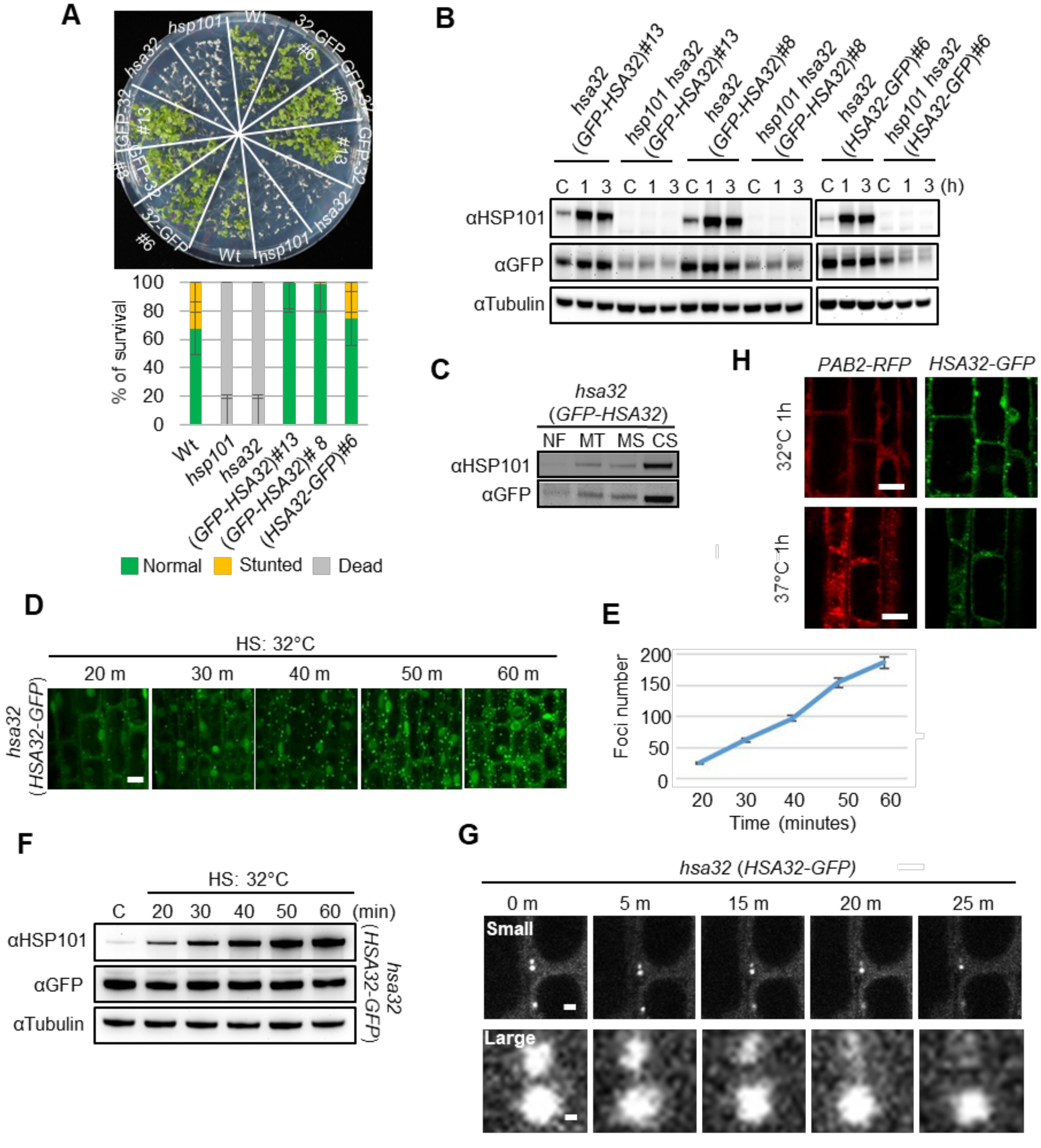


**Supplemental Figure S3. GFP-fused transgenic lines of HSA32. A)** Complementation analysis of GFP-fused HSA32 transgenic lines by using LAT assay. The transgenic lines with GFP-tagged HSA32 rescued the *hsa32* phenotype. **B)** HSA32 protein accumulation was examined in GFP-HSA32 and HSA32-GFP transgenic lines with *hsa32* and *hsp101 hsa32* background. **C)** Subcellular compartments of heat-acclimated seedlings were isolated from *hsa32* (*GFP-HSA32*) using differential ultra-centrifugation of their cell lysates. Four distinct fractions were collected and analyzed: nuclear fraction (NF), mitochondrial fraction (MF), microsomal fraction (MS), and cytosolic soluble fraction (CS). **D)** Time-lapse imaging of HSA32-GFP was performed to observe the foci formation following heat stress at 32°C for indicated time points. Scale bar, 20 µm. **E)** The graph represents the foci number against the periods of heat treatment at 32°C indicated in Figure D. **F)** HSA32 protein levels were examined using *hsa32* (*HSA32-GFP*) at the indicated time points. **G)** The upper and lower panels represented selected two foci of *hsa32* (*HSA32-GFP*) with small and large images to monitor whether the foci of HSA32 undergo a fusion event. Scale bars, 2 µm. **H)** The foci formation was observed in the transgenic line (*PAB2-RFP X HSA32-GFP*) at 32°C and 37°C HS conditions. Scale bars, 20 µm.

### Supplemental Figure S4

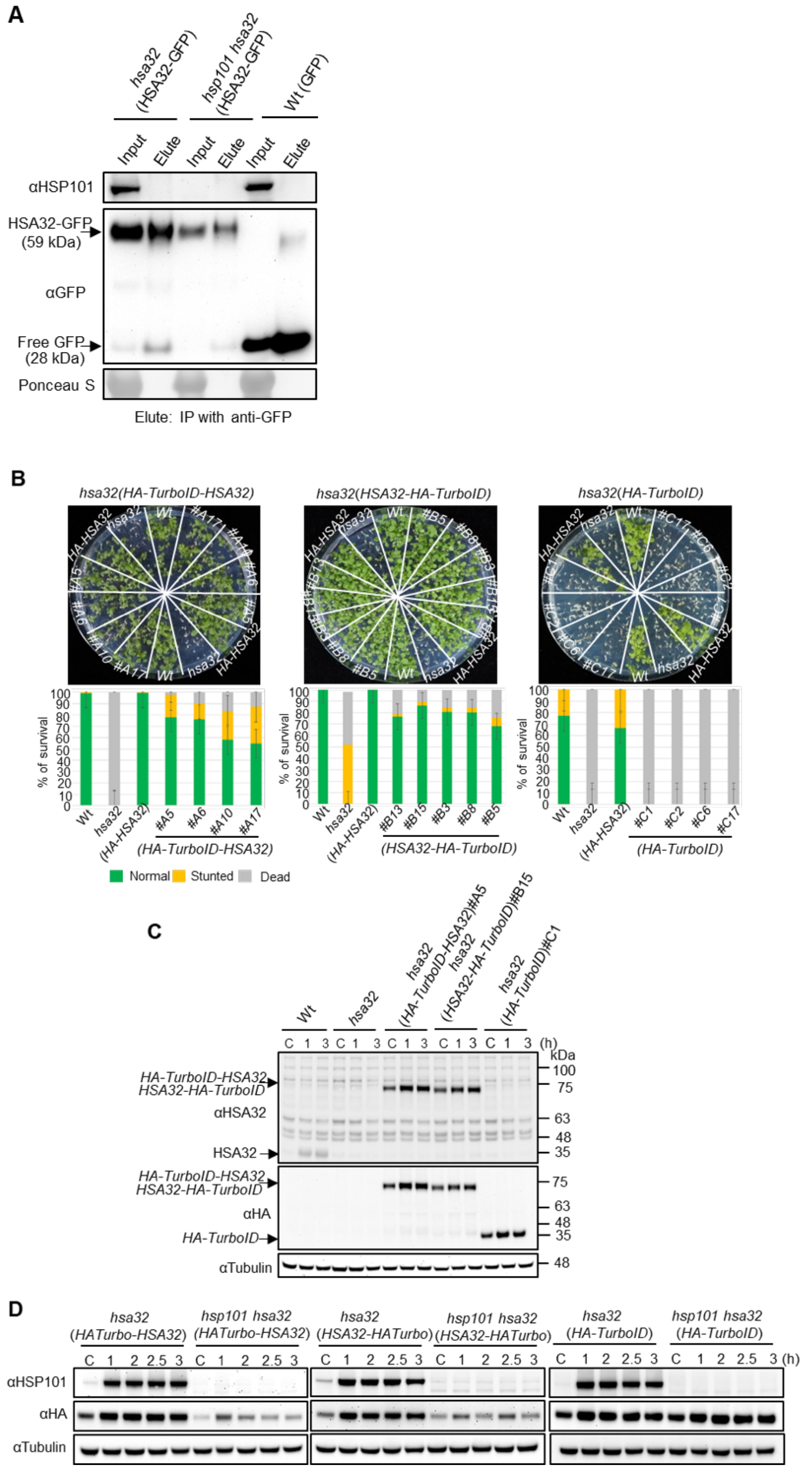

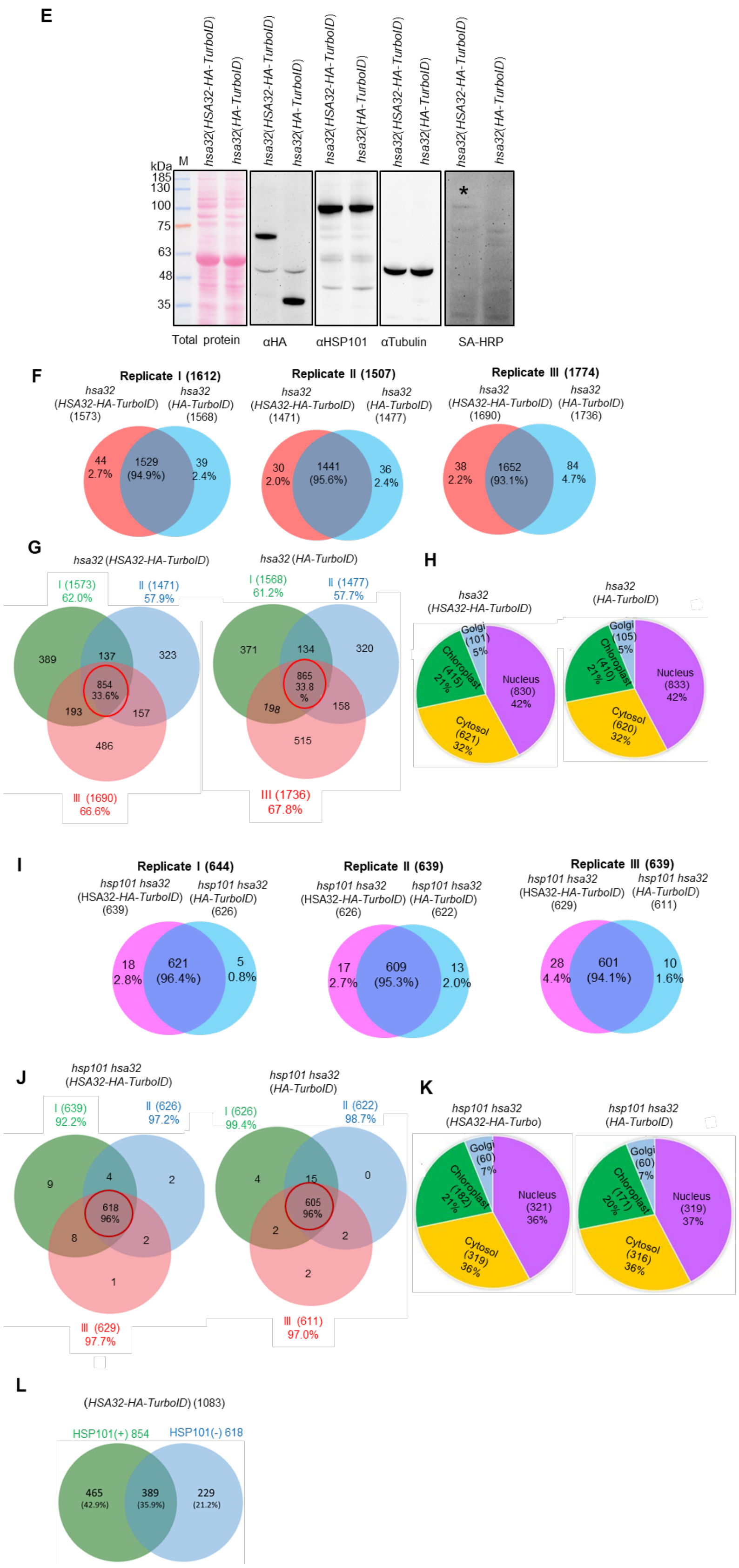
E
hsa32(HSA32-HA-TurboID)
hsa32(HA-TurboID)
kDa M
185
130
100
75
63
48
35
*
Total protein
αHA
αHSP101
αTubulin
SA-HRP
F
Replicate I (1612)
hsa32 (HSA32-HA-TurboID) (1573)
hsa32 (HA-TurboID) (1568)
44 2.7%
1529 (94.9%)
39 2.4%
Replicate II (1507)
hsa32 (HSA32-HA-TurboID) (1471)
hsa32 (HA-TurboID) (1477)
30 2.0%
1441 (95.6%)
36 2.4%
Replicate III (1774)
hsa32 (HSA32-HA-TurboID) (1690)
hsa32 (HA-TurboID) (1736)
38 2.2%
1652 (93.1%)
84 4.7%
G
hsa32 (HSA32-HA-TurboID)
I (1573) 62.0%
II (1471) 57.9%
389
137
323
854 33.6%
193
157
486
III (1690) 66.6%
hsa32 (HA-TurboID)
I (1568) 61.2%
II (1477) 57.7%
371
134
320
865 33.8 %
198
158
515
III (1736) 67.8%
H
hsa32 (HSA32-HA-TurboID)
Golgi (101) 5%
Chloroplast (415) 21%
Nucleus (830) 42%
Cytosol (621) 32%
hsa32 (HA-TurboID)
Golgi (105) 5%
Chloroplast (410) 21%
Nucleus (833) 42%
Cytosol (620) 32%
I
Replicate I (644)
hsp101 hsa32 (HSA32-HA-TurboID) (639)
hsp101 hsa32 (HA-TurboID) (626)
18 2.8%
621 (96.4%)
5 0.8%
Replicate II (639)
hsp101 hsa32 (HSA32-HA-TurboID) (626)
hsp101 hsa32 (HA-TurboID) (622)
17 2.7%
609 (95.3%)
13 2.0%
Replicate III (639)
hsp101 hsa32 (HSA32-HA-TurboID) (629)
hsp101 hsa32 (HA-TurboID) (611)
28 4.4%
601 (94.1%)
10 1.6%
J
hsp101 hsa32 (HSA32-HA-TurboID)
I (639) 92.2%
II (626) 97.2%
9
4
2
618 96%
8
2
1
III (629) 97.7%
hsp101 hsa32 (HA-TurboID)
I (626) 99.4%
II (622) 98.7%
4
15
0
605 96%
2
2
2
III (611) 97.0%
K
hsp101 hsa32 (HSA32-HA-Turbo)
Golgi (60) 7%
Chloroplast (182) 21%
Nucleus (321) 36%
Cytosol (319) 36%
hsp101 hsa32 (HA-TurboID)
Golgi (60) 7%
Chloroplast (171) 20%
Nucleus (319) 37%
Cytosol (316) 36%
L
(HSA32-HA-TurboID) (1083)
HSP101(+) 854
HSP101(-) 618
465 (42.9%)
389 (35.9%)
229 (21.2%)

**Supplemental Figure S4. Analysis of HSA32 interacting proteins by co-immunoprecipitation and proximity labeling. A)** Co-immunoprecipitation (Co-IP) assay was employed to detect the interaction between HSP101 and HSA32. We used transgenic plants expressing HSA32-GFP for the Co-IP assay to see if HSP101 can co-immunoprecipitate by GFP antibodies. Unfortunately, Co-IP was unable to capture their interaction. **B)** The LAT assay was conducted to analyze the complementation of the transgenic lines expressing HA-TurboID fused to the N- or C-terminus of HSA32 in the *hsa32* mutant background. HA-TurboID expressed in *hsa32* served as a negative control. Four idependent lines of *HA-TurboID-HSA32* (#A5, 6, 10, and 17), five independent lines of *HSA32-HA-TurboID* (#B3, 5, 8, 13 and15), and four independent lines of *HA-TurboID* (#C1, 2, 6, and 17) were used for complementation assay. **C)** The protein accumulation pattern of HA-TurboID was examined in the complementation lines following control and heat acclimation conditions using anti-HSA32 and HA antibodies. **D)** TurboID fused three complementation lines (N-and C-terminally fused HA-TurboID with HSA32, and HA-TurboID alone) were crossed with *hsp101 hsa32,* and homozygous lines were selected through genotyping. Immunoblotting was conducted to examine the protein level of HSA32 and HSP101 in all these transgenic lines. The blots were probed with anti-HSP101, HA, and tubulin antibodies. **E)** The biotinylated protein levels in *hsa32* (*HSA32-HA-TurboID*) *and hsa32* (*HA-TurboID*). Four-day-old seedlings were submerged in biotin solution (100 μM) for 3 h. Following protein extraction, excess biotin was removed using a PD-10 desalting column. For immunoblotting, SA-HRP was utilized to detect the biotinylated proteins, while an anti-HA antibody was employed to identify HA-TurboID expression and an HSP101 antibody was used to identify HSP101 protein in the immunoblotting analysis. Biotinylated bands close to 100 kDa were marked with asterisks. Tubulin was used as a loading control. The molecular weights (kDa) were labeled on the left. **F)** Venn diagram showing overlaps of the identified proteins between *HSA32-HA-TurboID* and *HA-TurboID* transgenic plants in *hsa32* background. Red and blue circles denoted the samples *hsa32 (HSA32-HA-TurboID)* and *hsa32* (*HA-TurboID*), respectively, in which *hsa32* (*HA-TurboID*) served as control. Three independent biological replicates were used in this Venn diagram. **G)** The left panel represents a Venn diagram illustrating the overlap in the number of proteins identified across the three biological replicates of *hsa32* (*HSA32-HA-TurboID*). The right panel depicts a Venn diagram visualizing the overlap in the number of proteins identified within the control samples *hsa32* (*HA-TurboID*) across three biological replicates. The green, blue, and pink circles represent replicates I, II, and III. **H)** The cellular distributions of all identified proteins of *hsa32* (*HSA32-HA-TurboID*) and *hsa32* (*HA-TurboID*) were visualized using pie charts. Here, the distribution of the nucleus, cytosol, chloroplast, and Golgi apparatus is denoted by purple, yellow, green, and blue color, respectively. **I)** Venn diagrams depict the overlap between the number of proteins identified in (*HSA32-HA-TurboID*) and (*HA-TurboID*) transgenic plants within a double mutant *hsa32 hsp101* background in three biological replicates. The red circle represents *hsp101 hsa32 (HSA32-HA-TurboID)* samples, while the blue circle denotes *hsp101 hsa32* (*HA-TurboID*) control samples. Data were obtained from three independent biological replicates. **J)** The left panel depicts a Venn diagram illustrating the overlap proteins identified across the three biological replicates of *hsp101 hsa32* (*HSA32-HA-TurboID*). The right panel represents a Venn diagram visualizing the overlap in the number of proteins identified within the control samples *hsp101 hsa32* (*HA-TurboID*) across three biological replicates. Green, blue, and pink circles denote I, II, and III replicates. **K)** The left and right panels depict pie charts showing the cellular component distribution of the identified proteins in *hsp101 hsa32* (*HSA32-HA-TurboID)* and *hsp101 hsa32* (*HA-TurboID*) samples, respectively. The distribution of the proteins localized in the nucleus, cytosol, chloroplast, and

Golgi apparatus is denoted by purple, yellow, green, and blue color, respectively. **L)** The Venn diagram showing the shared and unique proteins between the samples with (+) and without (-) HSP101 of *HSA32-HA-TurboID*.

**Supplemental Figure S5**

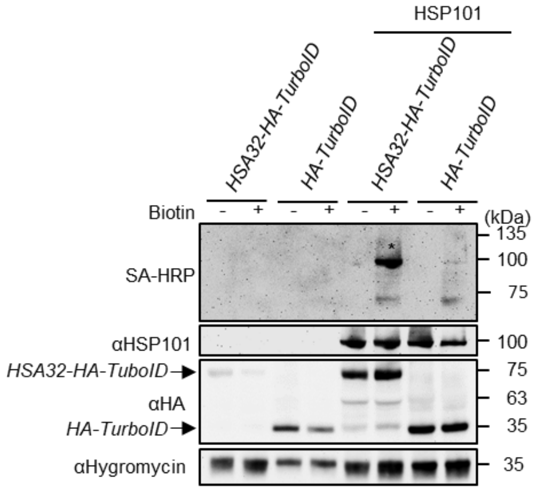


**Supplemental Figure S5. Agro-infiltration for transient co-expression of TurboID fused with HSA32 and Arabidopsis HSP101 in *N. benthamiana* leaves.** Agrobacterium strains containing the constructs for HSA32-HA-TurboID or HA-TurboID were co-infiltrated with or without the agrobacterium carrying the construct for HSP101. Co-infiltration was conducted in the leaves of three-week-old plants, followed by the infiltration of biotin (+) or buffer (-) into the same leaf sectors after agroinfiltration. The infiltrated leaves were collected for protein extraction one day after biotin infiltration. SA-HRP and antibodies against HSP101 and HA were used to probe the blots. The asterisks marked for the biotinylated proteins were approximately 100 kDa in size.

#### Supplemental Figure S6

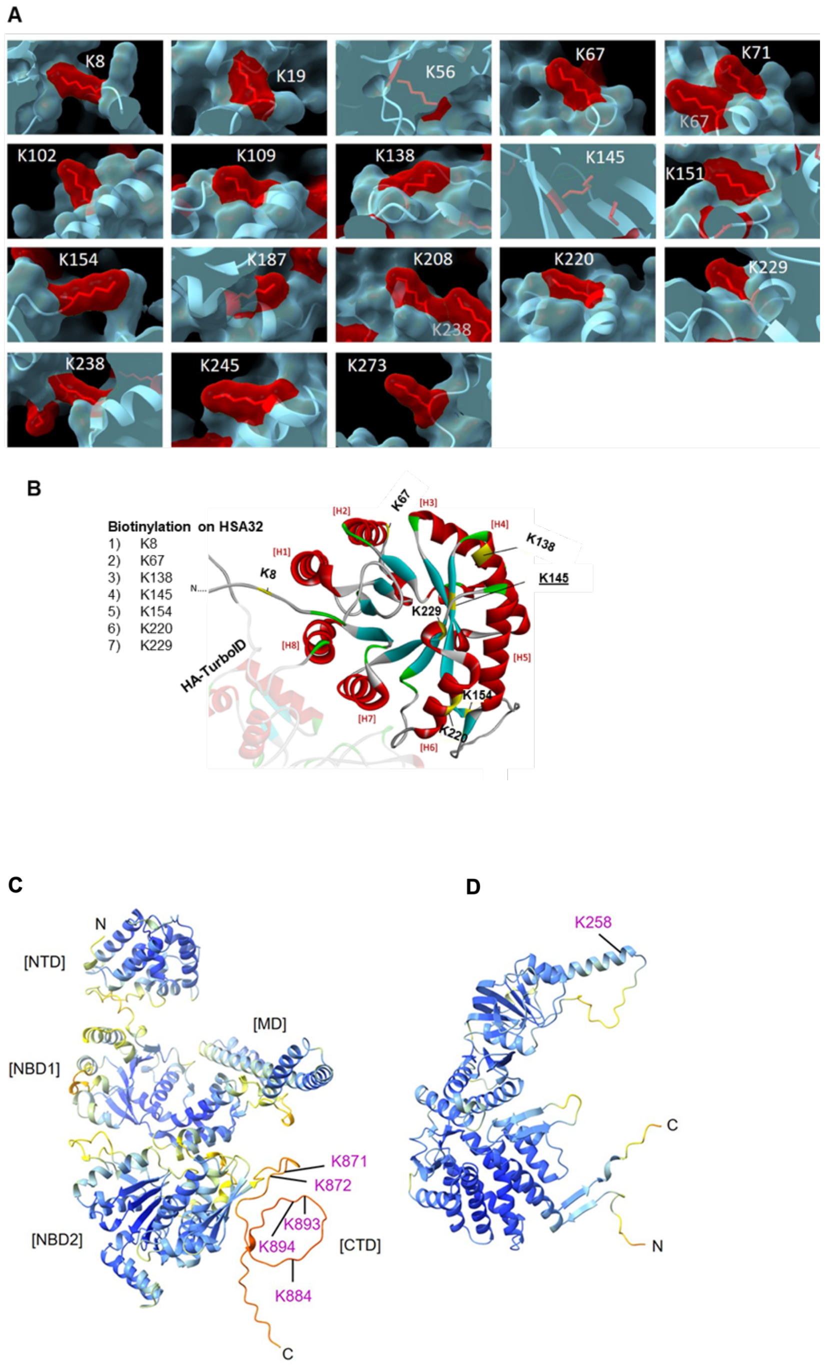


**Supplemental Figure S6. Biotinylation of lysine residues on HSA32-HA-TurboID and interacting proteins. A)** The protein structure of HSA32-HA-TurboID was predicted by AlphaFold 2. The molecular surfaces and atoms of 18 lysine residues on HSA32 were generated by ChimeraX and colored red. 16 out of 18 lysines of HSA32 were located on the surface of the predicted structure of HSA32-HA-TurboID, while there were two lysines, K56 and K145, embedded in the protein's interior. Due to the limitation of the mass spectrometer, the peptides containing K71, K109, or K187 could not be detected. In the predicted structure of the trimeric HSA32, the long loop would block the access of K208, K238, and K245 residues. These contribute to why biotinylation of some lysine residues could not be found. **B)** Seven biotinylated lysine (K) residues located on the 3-D structure of HSA32, predicted by AlphaFold2 software. Six biotinylated lysine residues were located on the surface of the 3-D structure of HSA32, while one lysine (K145) appeared to be embedded within the structure. **C)** Five biotinylated lysine residues were identified on the C-terminus of HSP101. **D)** One lysine residue (K258) on CCT2 was biotinylated. N: N-terminus; C: C-terminus: NTD: N-terminal domain; NBD1: Nucleotide-binding domain; CTD: C-terminal domain.

**Supplemental Figure S7**

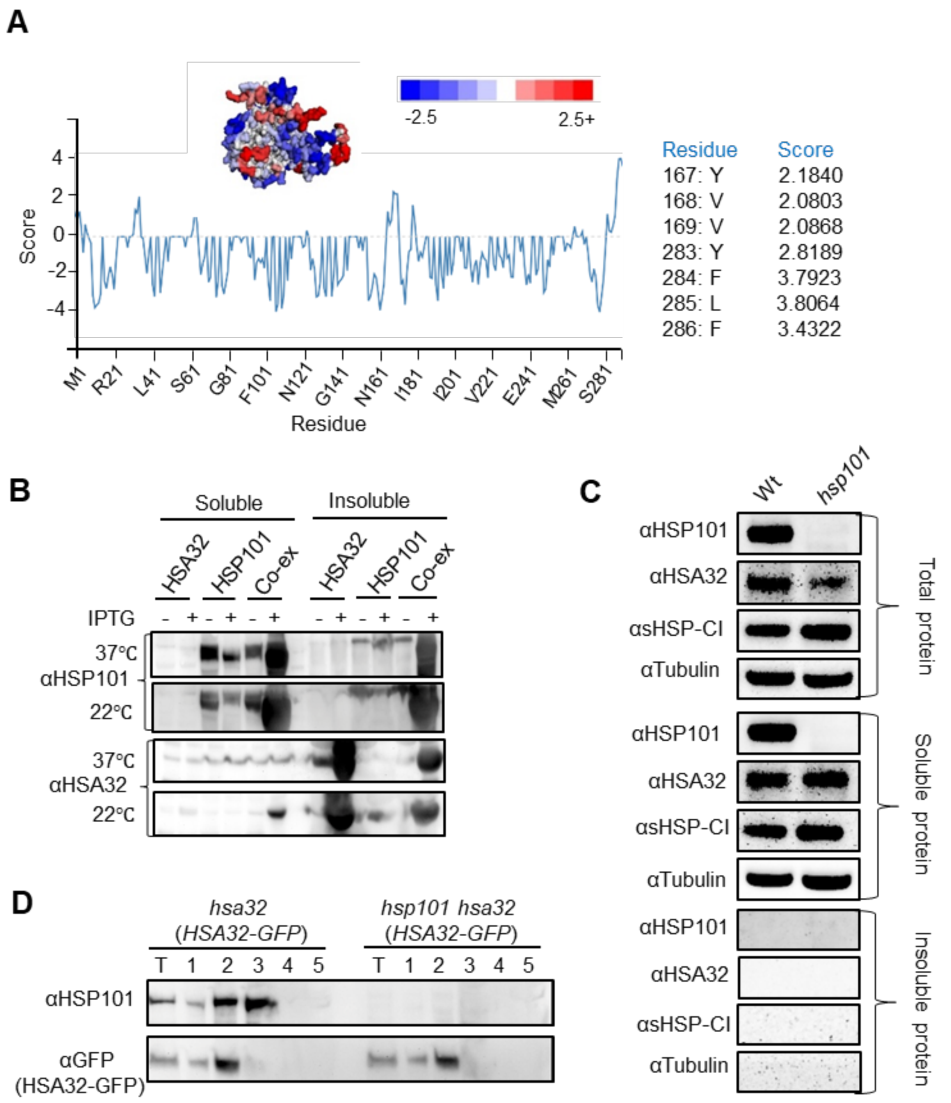


**Supplemental Figure S7. HSA32 is an aggregation-prone protein. A)** AGGRESCAN3D (A3D) was applied to predict the structural aggregation tendency of HSA32 based on AlphaFold2 predicted structure. The A3D structure of HSA32 displayed elevated A3D scores, with the red color indicating more aggregation-prone residues (Upper panel). A3D plot (lower panel) revealed notably high A3D scores (more than 2.0) in the amino acid residues 167-169 (loop) and 283-286 (C-terminal). **B)** The recombinant proteins, HSA32-His$_6$ and HSP101, were expressed in *E. coli*, both individually or together (Co-ex) under 22°C and 37°C conditions with or without IPTG induction. **C)** The solubility test of HSA32 was performed in Arabidopsis by following the protocol developed by McLoughlin et al. (2019). The seedlings of wild type and *hsp101* were subjected to heat treatment at 32°C for 1 h and collected samples during 1 h recovery period.

HSA32 protein remains in soluble fractions both in wild type and *hsp101* mutant. The soluble fractions represent the supernatant, whereas the insoluble fractions represent the repeatedly washed pellets resuspended in an equal buffer volume. The blots were probed with antibodies against HSP101, HSA32, sHSP-CI, and tubulin. In this experiment, HSP101, and sHSP-CI were identified in soluble fractions, but not insoluble fractions following heat acclimation which served as control. **D**) Ammonium sulfate precipitation of HSA32-GFP from the crude extract of Arabidopsis transgenic lines *hsa32* (*HSA32-GFP*) and *hsp101 hsa32* (*HSA32-GFP*). T, total protein; 1, 20~30% cut; 2, 30~40% cut; 3, 40~50% cut; 4, 50~60% cut; 5, 60~70% cut. The proteins were analyzed by SDS-PAGE and immunoblots with antibodies against HSP101 and GFP.
