## Supplemental Tables for "Stabilization of HSA32, an aggregation-prone protein, by the protein disaggregase HSP101 plays a critical role in maintaining acquired thermotolerance in Arabidopsis"

**Supplement Table S1: Cellular components of the identified proteins in *hsa32* background**

Significantly over-represented cellular components were set with FDR < 0.05. Only nucleus, cytosol, chloroplast, and Golgi apparatus are listed.
Fold enrichment = Number in Input List / Number in Reference

**Replicate I**

| *hsa32 (HSA32-HA-TurboID)* | |
|---|---|
| Uniquely Mapped IDs | 1555 out of **1562** |
| Unmapped IDs | 18 |
| Multiple mapping information | 5 |

| GO cellular component | Number in Input List | Number in Reference | Fold Enrichment | rawP value | FDR |
|---|---|---|---|---|---|
| **Nucleus** | **830** | 602.4 | **1.38** | 1.21E-29 | 4.98E-28 |
| **Cytosol** | **621** | 151.38 | **4.10** | 1.59E-199 | 1.57E-196 |
| **Chloroplast** | **415** | 287.45 | **1.44** | 1.54E-14 | 3.10E-13 |
| **Golgi apparatus** | **101** | 67.07 | **1.51** | 1.35E-04 | 1.14E-03 |

| *hsa32 (HA-TurboID)* | |
|---|---|
| Uniquely Mapped IDs | 1550 out of **1557** |
| Unmapped IDs | 11 |
| Multiple mapping information | 8 |

| GO cellular component | Number in Input List | Number in Reference | Fold Enrichment | rawP value | FDR |
|---|---|---|---|---|---|
| **Nucleus** | **833** | 600.47 | **1.39** | 5.95E-31 | 2.44E-29 |
| **Cytosol** | **620** | 150.9 | **4.11** | 1.19E-199 | 1.17E-196 |
| **Chloroplast** | **410** | 286.53 | **1.43** | 8.53-14 | 1.72E-12 |
| **Golgi apparatus** | **105** | 66.85 | **1.57** | 1.98E-05 | 2.01E-04 |

**Replicate II**

| *hsa32 (HSA32-HA-TurboID)* | |
|---|---|
| Uniquely Mapped IDs | 1450 out of **1452** |
| Unmapped IDs | 23 |
| Multiple mapping information | 19 |

| GO cellular component | Number in Input List | Number in Reference | Fold Enrichment | rawP value | FDR |
|---|---|---|---|---|---|
| **Nucleus** | **744** | 559.98 | **1.33** | 1.98E-21 | 4.55E-20 |
| **Cytosol** | **616** | 140.72 | **4.38** | 2.35E-215 | 2.32E-212 |
| **Chloroplast** | **399** | 267.21 | **1.49** | 1.81E-16 | 3.65E-15 |
| **Golgi apparatus** | **119** | 62.34 | **1.91** | 1.83E-10 | 3.23E-09 |

| *hsa32 (HA-TurboID)* | |
|---|---|
| Uniquely Mapped IDs | 1456 out of **1458** |
| Unmapped IDs | 22 |
| Multiple mapping information | 19 |

| GO cellular component | Number in Input List | Number in Reference | Fold Enrichment | rawP value | FDR |
|---|---|---|---|---|---|
| **Nucleus** | **747** | 562.29 | **1.33** | 1.44E-21 | 3.73E-20 |
| **Cytosol** | **626** | 141.3 | **4.43** | 7.24E-222 | 7.14E-219 |
| **Chloroplast** | **399** | 286.31 | **1.49** | 3.66E-16 | 7.37E-15 |
| **Golgi apparatus** | **117** | 62.6 | **1.87** | 1.01E-09 | 1.68E-08 |

**Replicate III**

| *hsa32 (HSA32-HA-TurboID)* | |
|---|---|
| Uniquely Mapped IDs | 1671 out of **1678** |
| Unmapped IDs | 20 |
| Multiple mapping information | 11 |

| GO cellular component | Number in Input List | Number in Reference | Fold Enrichment | rawP value | FDR |
|---|---|---|---|---|---|
| **Nucleus** | **847** | 647.14 | **1.31** | 8.78E-22 | 2.28E-20 |
| **Cytosol** | **659** | 162.63 | **4.05** | 3.61E-207 | 3.55E-204 |
| **Chloroplast** | **468** | 308.8 | **1.52** | 4.91E-20 | 1.18E-18 |
| **Golgi apparatus** | **128** | 72.05 | **1.78** | 3.98E-09 | 5.31E-08 |

| *hsa32 (HA-TurboID)* | |
|---|---|
| Uniquely Mapped IDs | 1456 out of **1458** |
| Unmapped IDs | 22 |
| Multiple mapping information | 19 |

| GO cellular component | Number in Input List | Number in Reference | Fold Enrichment | rawP value | FDR |
|---|---|---|---|---|---|
| **Nucleus** | **874** | 664.49 | **1.32** | 3.88E-23 | 1.01E-21 |
| **Cytosol** | **673** | 166.99 | **4.03** | 1.30E-209 | 1.29E-206 |
| **Chloroplast** | **473** | 317.08 | **1.49** | 6.16E-19 | 1.48E-17 |
| **Golgi apparatus** | **129** | 73.98 | **1.74** | 9.12E-09 | 1.20E-07 |

**Supplement Table S2: HSA32 interacting proteins identified by TurboID proximity labeling**

All the HSA32 interacting proteins matched the criteria, namely the abundance ratio of the identified protein in *hsa32 (HSA32-HA-TruboID)* versus *hsa32 (HA-TurboID)* $\geq 2$ and the *p* value < 0.05 in three biological replicates. PSM: peptide-spectrum match

| No | Gene Name / Accession | Protein Name | Coverage (%) | #PSM | #Peptide | # Unique peptide | Abundance ratio | *p* value |
|---|---|---|---|---|---|---|---|---|
| 1 | *HSA32* / AT4G21320 | Heat stress-associated 32-kD protein | 57.0 | 212 | 24 | 24 | — | — |
| 2 | *HSP101* / AT1G74310 | Heat shock protein 101 | 72.9 | 2142 | 107 | 101 | 30.0 | 4.36E-04 |
| 3 | *CCT8* / AT3G03960 | Chaperonin CCT8 | 61.2 | 751 | 37 | 37 | 8.1 | 4.37E-03 |
| 4 | *ATJ3* / AT3G44110 | DNAJ homologue 3 ATJ3 (co-chaperone of HSP70) | 24.5 | 69 | 12 | 4 | 4.8 | 4.28E-03 |
| 5 | *CCT3* / AT5G26360 | Chaperonin CCT3 | 32.4 | 111 | 19 | 19 | 4.0 | 2.78E-02 |
| 6 | *P23-1* / AT4G02450 | P23-1 (co-chaperon of HSP90) | 43.6 | 233 | 14 | 14 | 3.5 | 3.84E-06 |
| 7 | *CCT2* / AT5G20890 | Chaperonin CCT2 | 34.7 | 106 | 17 | 17 | 2.1 | 1.03E-02 |

**Supplement Table S3: Cellular components of the identified proteins in *hsp101 hsa32* background**

Significantly over-represented cellular components were set with FDR < 0.05. Only nucleus, cytosol, chloroplast, and Golgi apparatus are listed.

Fold enrichment = Number in Input List / Number in Reference

**Replicate 1**

| *hsa32 hsp101 (HSA32-HA-TurboID)* | |
|---|---|
| Uniquely Mapped IDs | 631 out of **634** |
| Unmapped IDs | 8 |
| Multiple mapping information | 4 |

| GO cellular component | Number in Input List | Number in Reference | Fold Enrichment | rawP value | FDR |
|---|---|---|---|---|---|
| **Nucleus** | **321** | 243.82 | **1.32** | 8.79E-10 | 1.70E-08 |
| **Cytosol** | **319** | 61.26 | **5.21** | 3.08E-143 | 3.04E-140 |
| **Chloroplast** | **182** | 116.49 | **1.56** | 4.36E-10 | 8.77E-09 |
| **Golgi apparatus** | **60** | 27.11 | **2.21** | 3.47E-08 | 6.33E-07 |

| *hsa32 hsp101 (HA-TurboID)* | |
|---|---|
| Uniquely Mapped IDs | 618 out of **620** |
| Unmapped IDs | 8 |
| Multiple mapping information | 3 |

| GO cellular component | Number in Input List | Number in Reference | Fold Enrichment | rawP value | FDR |
|---|---|---|---|---|---|
| **Nucleus** | **319** | 238.43 | **1.34** | 1.09E-10 | 2.23E-09 |
| **Cytosol** | **316** | 59.91 | **5.27** | 4.94E-144 | 4.88E-141 |
| **Chloroplast** | **171** | 113.92 | **1.50** | 3.47E-08 | 6.46E-07 |
| **Golgi apparatus** | **60** | 26.51 | **2.26** | 1.31E-08 | 2.48E-07 |

**Replicate 2**

| *hsa32 hsp101 (HSA32-HA-TurboID)* | |
|---|---|
| Uniquely Mapped IDs | 618 out of **621** |
| Unmapped IDs | 8 |
| Multiple mapping information | 4 |

| GO cellular component | Number in Input List | Number in Reference | Fold Enrichment | rawP value | FDR |
|---|---|---|---|---|---|
| **Nucleus** | **316** | 238.82 | **1.32** | 5.87E-10 | 1.14E-08 |
| **Cytosol** | **316** | 60.00 | **5.27** | 9.01E-144 | 8.88E-141 |
| **Chloroplast** | **180** | 114.1 | **1.58** | 2.08E-10 | 4.10E-09 |
| **Golgi apparatus** | **60** | 26.55 | **2.26** | 1.36E-08 | 2.49E-07 |

| *hsa32 hsp101 (HA-TurboID)* | |
|---|---|
| Uniquely Mapped IDs | 614 out of **616** |
| Unmapped IDs | 8 |
| Multiple mapping information | 3 |

| GO cellular component | Number in Input List | Number in Reference | Fold Enrichment | rawP value | FDR |
|---|---|---|---|---|---|
| **Nucleus** | **318** | 236.89 | **1.34** | 7.13E-11 | 1.47E-09 |
| **Cytosol** | **316** | 59.52 | **5.31** | 4.42E-145 | 4.36E-142 |
| **Chloroplast** | **169** | 113.18 | **1.49** | 5.72E-08 | 1.06E-06 |
| **Golgi apparatus** | **60** | 26.34 | **2.28** | 1.14E-08 | 2.16E-07 |

**Replicate 3**

| *hsa32 hsp101 (HSA32-HA-TurboID)* | |
|---|---|
| Uniquely Mapped IDs | 621 out of **624** |
| Unmapped IDs | 8 |
| Multiple mapping information | 4 |

| GO cellular component | Number in Input List | Number in Reference | Fold Enrichment | rawP value | FDR |
|---|---|---|---|---|---|
| **Nucleus** | **317** | 239.97 | **1.32** | 6.56E-10 | 1.27E-08 |
| **Cytosol** | **317** | 60.29 | **5.26** | 6.19E-144 | 6.10E-141 |
| **Chloroplast** | **179** | 114.65 | **1.56** | 6.23E-10 | 1.23E-08 |
| **Golgi apparatus** | **60** | 26.68 | **2.25** | 1.54E-08 | 2.82E-07 |

| *hsa32 hsp101 (HA-TurboID)* | |
|---|---|
| Uniquely Mapped IDs | 603 out of **605** |
| Unmapped IDs | 8 |
| Multiple mapping information | 3 |

| GO cellular component | Number in Input List | Number in Reference | Fold Enrichment | rawP value | FDR |
|---|---|---|---|---|---|
| **Nucleus** | **318** | 232.66 | **1.37** | 4.51E-12 | 9.26E-11 |
| **Cytosol** | **314** | 58.46 | **5.37** | 4.22E-146 | 4.16E-143 |
| **Chloroplast** | **166** | 111.16 | **1.49** | 7.89E-08 | 1.47E-06 |
| **Golgi apparatus** | **58** | 25.87 | **2.24** | 2.97E-08 | 5.64E-07 |

**Supplement Table S4: HSA32 interacting proteins identified by TurboID proximity labeling**

All the HSA32 interacting proteins matched the criteria, namely the abundance ratio of the identified protein in *hsp101 hsa32 (HSA32-HA-TruboID)* versus *hsp101 hsa32 (HA-TurboID) ≥ 2* and the *p* value < 0.05 in three biological replicates.

PSM: peptide-spectrum match

| No | Gene Name / Accession | Protein Name | Coverage (%) | #PSM | #Peptide | # Unique peptide | Abundance ratio | *p* value |
|---|---|---|---|---|---|---|---|---|
| 1 | *HSA32* / AT4G21320 | Heat stress-associated 32-kD protein | 48 | 125 | 17 | 17 | — | — |
| 2 | *CCT3* / AT5G26360 | Chaperonin CCT3 | 23 | 58 | 11 | 11 | 29.9 | 8.54E-04 |
| 3 | *CCT8* / AT3G03960 | Chaperonin CCT8 | 55 | 374 | 30 | 30 | 13.7 | 2.07E-03 |
| 4 | *ATJ3* / AT3G44110 | Chaperone protein dnaJ 3 | 10 | 8 | 4 | 4 | 5.7 | 7.41E-03 |
| 5 | *CCT6A/B* / AT3G02530 AT5G16070 | Chaperonin CCT6A/B | 16 | 39 | 8 | 8 | 5.4 | 9.32E-03 |
| 6 | *CCT2* / AT5G20890 | Chaperonin CCT2 | 13 | 28 | 6 | 6 | 4.0 | 2.10E-02 |
| 7 | *P23-1* / AT4G02450 | Co-chaperone protein p23-1 | 33 | 104 | 8 | 9 | 3.9 | 1.57E-02 |
| 8 | *IPA1* / AT1G18660 | Zinc finger (C3HC4-type RING finger) family protein | 24 | 92 | 14 | 14 | 3.5 | 2.53E-02 |
| 9 | *TUBA3* / AT5G19770 | Tubulin alpha-3 chain | 8 | 26 | 3 | 1 | 3.2 | 4.30E-02 |
| 10 | *TUBA6* / AT4G14960 | Tubulin alpha-6 chain | 26 | 88 | 8 | 6 | 3.1 | 3.46E-02 |
| 11 | *TUBB5* / AT1G20010 | Tubulin beta-5 chain | 8 | 17 | 3 | 1 | 2.6 | 3.75E-02 |
| 12 | *BT4* / AT5G67480 | BTB and TAZ domain protein 4 | 2 | 1 | 1 | 1 | 2.3 | 4.09E-02 |
| 13 | *LEA25* / AT2G42560 | Late embryogenesis abundant domain protein | 52 | 342 | 30 | 30 | 2.2 | 4.93E-02 |

**Supplemental Table S5: Combinations of different volumes (mL) of agrobacteria for modulating the expression levels of HSP101, HA-HSP101, HSP101-HA, and HSA32-HA-TurboID in Figure 5E**

| HSA32-TurboID | 0.1 | 0.1 | 0.2 | 0.2 | 1 | 1 | 1 | 0.1 | 0.2 | 0.3 |
|---|---|---|---|---|---|---|---|---|---|---|
| HSP101 | 0.2 | 0.3 | 0.2 | 0.3 | 0 | 0 | 0 | 0 | 0 | 0 |
| HA-HSP101 | 0 | 0 | 0 | 0 | 0.5 | 0.6 | 0.7 | 0 | 0 | 0 |
| HSP101-HA | 0 | 0 | 0 | 0 | 0 | 0 | 0 | 0.9 | 0.9 | 0.9 |
| Suspension solution | 1.7 | 1.6 | 1.6 | 1.5 | 0.5 | 0.4 | 0.3 | 1 | 0.9 | 0.8 |
| Total | | | | | 2 | | | | | |

### Supplemental Table S6: Identified unique peptides and biotinylation sites on HSA32-HA-TurboID

| Positions | Annotated Sequence | # PSMs | Found in Samples | | Abundances | |
|---|---|---|---|---|---|---|
| | | | HSP101(+) | HSP101(-) | HSP101(+) | HSP101(-) |
| [2-8] | [M].A$^{Ac}$AYYRWK | 5 | | | 0 | 3,938,424 |
| [2-19] | [M].A$^{Ac}$AYYRW**K**$^{Bio}$SFEENEDRPEK **(K8)** | 1 | | | 422,036 | 0 |
| [7-19] | [R].W**K**$^{Bio}$SFEENEDRPEK **(K8)** | 2 | | | 2,841,028 | 0 |
| [7-21] | [R].W**K**$^{Bio}$SFEENEDRPEKPR **(K8)** | 8 | | | 4,408,494 | 0 |
| [9-19] | [K].SFEENEDRPEK | 9 | | | 16,613,434 | 8,449,227 |
| [9-21] | [K].**SFEENEDRPEKPR** | 34 | | | **58,403,633** | **1,353,060** |
| [9-22] | [K].SFEENEDRPEKPRR | 1 | | | 900,846 | 0 |
| [22-29] | [R].RYGVTEMR | 2 | | | 25,083,572 | 0 |
| [23-29] | [R].YGVTEMR | 9 | | | 31,376,730 | 0 |
| [57-67] | [K].**FSGGSNSLIPK** | 32 | | | **66,079,834** | **13,190,134** |
| [57-71] | [K].FSGGSNSLIP**K**$^{Bio}$SFIK **(K67)** | 5 | | | 2,552,785 | 503,848 |
| [72-95] | [K].QAIEMAHEHGVYVSTGDWAEHMLR | 15 | | | 4,626,713 | 629,676 |
| [72-102] | [K].QAIEMAHEHGVYVSTGDWAEHMLRSGPSAFK | 2 | | | — | 0 |
| [136-143] | [R].LI**K**$^{Bio}$NGGLR **(K138)** | 2 | | | 824,214 | 0 |
| [144-151] | [R].A**K**$^{Bio}$PMFAVK **(K145)** | 4 | | | 8,629,657 | 0 |
| [152-160] | [K].FN**K**$^{Bio}$SDIPGR **(K154)** | 12 | | | 6,485,212 | 420,168 |
| [161-173] | [R].NRAFGSYVVPEPR | 3 | | | 198,661 | 0 |
| [163-173] | [R].AFGSYVVPEPR | 24 | | | 126,651,956 | 4,844,711 |
| [174-187] | [R].SSEFVEDIDLLIRK | 18 | | | 6,688,213 | 4,527,027 |
| [209-214] | [K].YADSLR | 9 | | | 5,365,120 | 2,146,854 |
| [215-224] | [R].ADIIA**K**$^{Bio}$VIGR **(K220)** | 12 | | | 5,643,639 | 574,719 |
| [221-229] | [K].VIGRLGIEK | 5 | | | 476,708 | 1,119,418 |
| [221-238] | [K].VIGRLGIE**K**$^{Bio}$TMFEASDAK **(K229)** | 9 | | | 0 | 1,107,694 |
| [225-238] | [R].LGIE**K**$^{Bio}$TMFEASDAK **(K229)** | 20 | | | 15,816,804 | 1,302,454 |
| [230-238] | [K].TMFEASDAK | 15 | | | 20,130,006 | 15,628 |
| [239-245] | [K].**LVEWFIK** | 24 | | | **85,103,504** | **22,772,838** |
